# PRMT1 is a critical dependency in clear cell renal cell carcinoma through its role in post-transcriptional regulation of DNA damage response genes

**DOI:** 10.1101/2023.03.30.534936

**Authors:** Joseph Walton, Angel SN Ng, Karen Arevalo, Anthony Apostoli, Jalna Meens, Christina Karamboulas, Jonathan St-Germain, Panagiotis Prinos, Julia Dmytryshyn, Eric Chen, Cheryl H. Arrowsmith, Brian Raught, Laurie Ailles

**Affiliations:** Department of Medical Biophysics, University of Toronto, Toronto, Ontario, Canada; Princess Margaret Cancer Centre, University Health Network, Toronto, Ontario, Canada; Structural Genomics Consortium, University of Toronto, Toronto, Ontario, Canada

## Abstract

Biallelic inactivation of the Von Hippel-Lindau (*VHL*) tumor suppressor gene occurs in almost all cases of clear cell renal cell carcinoma (ccRCC) and leads to disrupted oxygen sensing and the upregulation of hypoxia-related genetic programs. Although the loss of VHL appears to be a necessary oncogenic driver event in the majority of ccRCC instances, it is not always a sufficient one. Large genomics studies have revealed that co-deletions of *VHL* with genes involved in chromatin regulation are common and important co-drivers of tumorigenesis. Several conserved evolutionary subtypes have been described clinically and the majority implicate disruptions in epigenetic regulators. It is now clear that impairments in cellular epigenetic mechanisms are important co-drivers of disease and signal a potential vulnerability in the epigenetic network of ccRCC cells relative to their normal counterparts. Using a clinically relevant panel of ccRCC models, we herein sought to exploit this potential vulnerability by screening a library of small molecule inhibitors that target a spectrum of epigenetic regulators. We identified MS023, an inhibitor of type I protein arginine methyltransferases (PRMTs) as an agent with antitumor activity. Using orthogonal genetic technologies, we further validated PRMT1 as the specific critical dependency for cancer growth. Mechanistically, our transcriptomic and functional analyses suggest that MS023 treatment results in substantial impairments to cell cycle and DNA damage repair (DDR) pathways, while spawning an accumulation of DNA damage over time. Our PRMT1 specific proteomics analysis revealed an interactome rich in RNA binding proteins including the specific regulator of DDR mRNA metabolism: the BCLAF1/THRAP3 complex. Further investigation suggests that MS023 treatment may result in impairments to DDR specific mRNA activities including nucleocytoplasmic transport and RNA splicing. Together, our data supports PRMT1 as a compelling target in ccRCC and informs a potential mechanism-based strategy for translational development.

## Introduction

The clear cell subtype of renal cell carcinoma (ccRCC) is the most common malignancy to arise in the kidney and accounts for the majority of renal cancer related deaths^1^. If detected early, localized tumors are surgically resected with intent to cure^2^. However, many patients present with disseminated disease at the time of diagnosis^3^, and even with surgery disease recurrence is common^4^. The prognosis for metastatic ccRCC is poor. Five-year survival rates hover near 12%^5^ and, thanks to an insidious resistance to both radiation and cytotoxic chemotherapeutic agents, treatment options remain limited. It has been known for some time that biallelic inactivation of the *von Hippel Lindau* gene (*VHL*) occurs in most cases of ccRCC, leading to complete loss of pVHL and a constitutive activation of the cell’s hypoxia response^6^. Although this appears to be a necessary and near universal driver event, evidence suggests that it alone is insufficient to initiate tumorigenesis^7–9^. Recent, large-scale genomic analyses have revealed additional, frequent loss-of-function mutations in several key genes including *PBRM1, SETD2, BAP1*, *KDM5C, KDM6A, MLL2, ARID1A* and *ARID1B*^10, 11^. There exists a surprising and common mechanistic theme to these additional putative tumor suppressor genes (TSGs): their protein products are all involved in epigenomic regulation.

Given the frequency of gene mutations impacting epigenetic regulatory proteins, and considering recent evidence from genetically engineered mouse models (GEMMs) that substantiate the importance of epigenetic regulators like PBRM1 and BAP1 in ccRCC tumor establishment^12, 13^, we sought to identify epigenetic vulnerabilities that may be exploited to develop new therapies. To accomplish this, we performed an *in vitro* proliferative screen across a panel of patient derived ccRCC models^14^. We used a library of validated chemical probes^15, 16^ that selectively target a spectrum of epigenetic regulatory proteins. The results of our screen identified some previously characterized ccRCC targets, including the enhancer of zeste homolog 2 (EZH2), as well as a novel regulator of ccRCC growth: protein arginine methyltransferases (PRMTs).

The methylation of arginine residues on histones and non-histone proteins is a prevalent post translational modification (PTM) and important regulator of multiple cellular processes^17^. PRMTs are the only known family of enzymes responsible for catalyzing the transfer of methyl groups from the S-adenosylmethionine (SAM) cofactor to the terminal nitrogen atoms of the guanidino group of arginine residues^18^. All nine identified PRMTs catalyze the transfer of one methyl group to produce mono-methylarginine (MMA), and are further classified according to the final methylarginine species they generate: type I members transfer an additional group to the same guanidino nitrogen producing asymmetric dimethylarginine (aDMA), while type II members transfer an additional group to the other guanidino nitrogen to make symmetric dimethylarginine (sDMA), and the sole type III enzyme, PRMT7, only catalyzes the formation of MMA^18^. Although arginine methylation is a relatively understudied PTM, emerging evidence has linked the activity of PRMTs to a number of cellular mechanisms important for the development and growth of cancer including epigenetic-mediated gene expression, RNA metabolism, DNA damage response, stem cell function, and the immune response^19^. Accordingly, the PRMT family has garnered significant attention as a potential therapeutic target and clinical evaluation of type I PRMT inhibitors and PRMT5 selective inhibitors are underway^20, 21^.

In this study, we identified inhibition of type I PRMTs as a novel vulnerability in ccRCC and using orthogonal genetic approaches, we validated PRMT1 as the specific type I enzyme mediating growth arrest. PRMT1 is the canonical member of the type I family responsible for the majority of all aDMA species produced in the cell^18^. As such, we employed transcriptomic and proteomic approaches to investigate its role as a mediator of ccRCC growth and survival. Our data suggests that type I PRMT inhibition leads to a pronounced down regulation of the cell cycle, compromised DNA damage repair (DDR) pathways and an accumulation of double-strand breaks (DSBs). Consistent with other reports, proteomic analysis confirms PRMT1’s central role as a regulator of proteins involved in RNA metabolism^18, 22, 23^ including targets connected to the mRNA splicing and export of important DDR enzyme transcripts. Using patient derived xenografts (PDXs), we further validated the inhibition of type I PRMTs and PRMT1 specifically as a viable *in vivo* strategy for attenuating ccRCC tumor growth. Together, our data argues for the potential translational benefit of PRMT1 inhibition as a clinical therapeutic strategy in ccRCC.

## Results

### *In vitro* chemical probe screen identifies Type I PRMTs as mediators of ccRCC cell proliferation

Our lab has previously developed an efficient methodology for establishing patient-derived cell line models of ccRCC from primary tumor tissues^14^. To ensure faithful representation of the conserved evolutionary subtypes seen clinically^9^, a panel of seven patient-derived models and one commercially available and widely used ccRCC cell line (786-0) were selected to facilitate the chemical probe screen (Figure 1A). These models underwent targeted sequencing to confirm the presence of relevant evolutionary subtype epigenetic driver mutations including *VHL*, *PBRM1*, *SETD2*, and *BAP1*. Using a cell permeable, far-red DNA fluorescent dye (DRAQ5^TM^) and the LI-COR imaging system, growth of these models was interrogated after seven days exposure to a collection of 36 epigenetic chemical probes from the Structural Genomics Consortium (SGC: https://www.thesgc.org/chemical-probes/epigenetics)^15, 16^. This library of small molecule ‘epiprobes’ selectively targets key epigenetic regulatory proteins, including several targets with compounds currently in clinical development. The results of our screen (Figure 1B) identified a total of 10 compounds that significantly reduce cell proliferation across the aggregate of all cell line models tested by a minimum of 50% (Supplementary Figure 1). Included in this list was UNC1999, an inhibitor of the Polycomb group (PcG), H3K27me3 lysine methyltransferase EZH2, which has been previously characterized as a target in ccRCC^24–26^. Additionally, GSKJ4, an inhibitor of the lysine demethylases that act on methylated H3K27 (KDM6A and KDM6B) also registered as a hit in our screen, underscoring the importance of the H3K27me3 epigenetic mark in maintaining ccRCC growth. Intriguingly, MS023, an inhibitor of all type I PRMTs, significantly repressed cell proliferation while the more specific type I inhibitors, MS049 (targeting PRMT4 (CARM1) & PRMT6), and SGC707 (targeting PRMT3) did not. The type II PRMT inhibitor GSK591 (targeting PRMT5) also demonstrated significant growth inhibitory effects across our disease models. Follow up dose response experiments comparing the inhibition of type I PRMTs by MS023 to the activity of a chemically similar but inactive compound, MS094, demonstrated high specificity and potency for type I PRMT inhibition (Figure 1C and Supplemental Figure 2). MS023 IC_50_ values ranged from 0.4 µM to 6 µM across our cell line panel.

**Figure 1:**
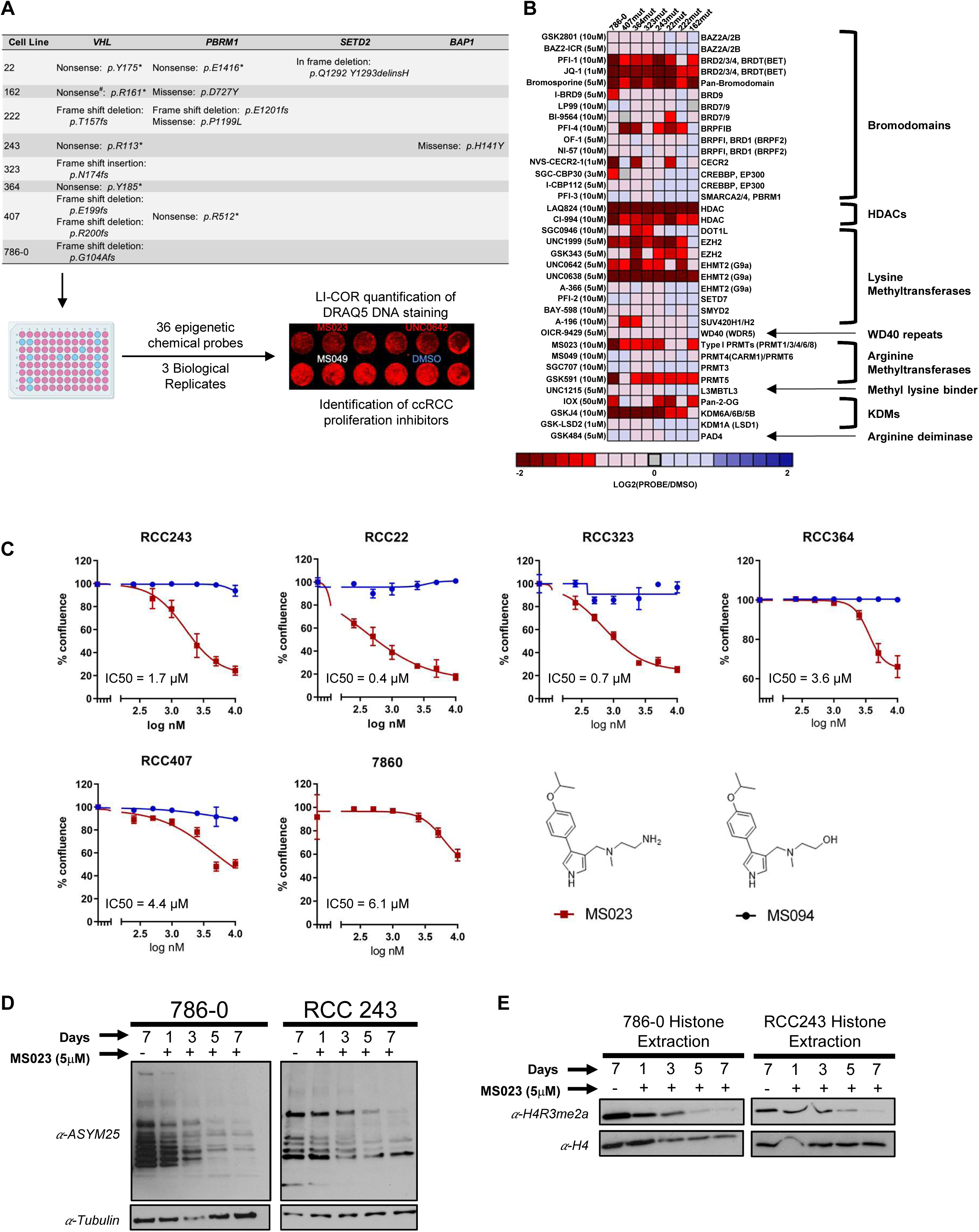
An epigenetic chemical probe screen identifies type I PRMT inhibitor MS023 as an inhibitor of ccRCC cell proliferation. **a,** Mutation profiles for select genes in eight ccRCC cell lines and the schematic work flow of epigenetic chemical probe screen. **b,** Heat map showing the average cell proliferation values in the presence of the indicated epigenetic chemical probe after seven days exposure in eight ccRCC cell lines (data shown as mean of n = 3). **c,** Dose response curves and MS023 IC50 values across a number of ccRCC models. Data are presented as the mean ± SEM calculated from at least 3 replicates for each cell line. Data presented at Day 5 for 786-0, Day 7 for RCC243, Day 8 for RCC407, Day 12 for RCC22 and Day 14 for RCC323 (determined based on the time at which control-treated cells reached confluence) **d,** Western blot analysis of asymmetric di-methylarginine (ADMA) changes in 786-0 and RCC243 cells after MS023 treatment. **e,** Western blot analysis of H4R3me2a changes in acid histone extractions of 786-0 and RCC243 following MS023 treatment.

To validate MS023’s on-target activity, we treated cell lines RCC243 and 786-0 for a period of 7 days with 5µM MS023 and using an aDMA specific antibody (ASYM25), we observed a global down regulation of aDMA species over time (Figure 1D). Additionally, since MS023 shows high potency against PRMT1 activity^27^, the type I enzyme solely responsible for the histone mark H4R3me2a^28^, we also used this post translational modification (PTM) as a proxy for PRMT1 inhibition. MS023 treatment followed by histone extraction and immunoblotting with an H4R3me2a-specific antibody resulted in a noticeable depletion of the PTM in both RCC243 and 786-0 (Figure 1E).

While type I PRMT inhibition has recently been attracting attention as a potential therapeutic tool in cancer treatment^21, 29^, to our knowledge only one study to date has suggested a role for PRMTs in ccRCC^30^, and no in-depth mechanistic investigations have been reported for these targets in kidney cancer. As such, we selected MS023 as our lead compound and sought to further characterize type I PRMT inhibition in ccRCC.

### The PRMT1 enzyme is the critical dependency among type I PRMTs in ccRCC

MS023 is known to have specific activity against all type I PRMT enzymes including PRMT1, PRMT3, PRMT4(CARM1), PRMT6 and PRMT8^27^. However, no decrease in cell proliferation was noted in our screen for the compounds MS049 and SGC707 that specifically target PRMT4 (CARM1)/PRMT6 and PRMT8, respectively. Transcription levels for each type I enzyme were compared across our previously sequenced cell line models^14^ and PRMT1 was found to have the highest relative expression (Figure 2A). Conversely, transcripts for PRMT8, a known membrane bound, neuronal specific protein^31^, were very low to undetectable. To confirm the functional importance of each type I PRMT enzyme for proliferation and viability, we employed a CRISPR-Cas9, GFP-drop out methodology (adapted from Lin *et al*^32^; Figure 2B).

**Figure 2:**
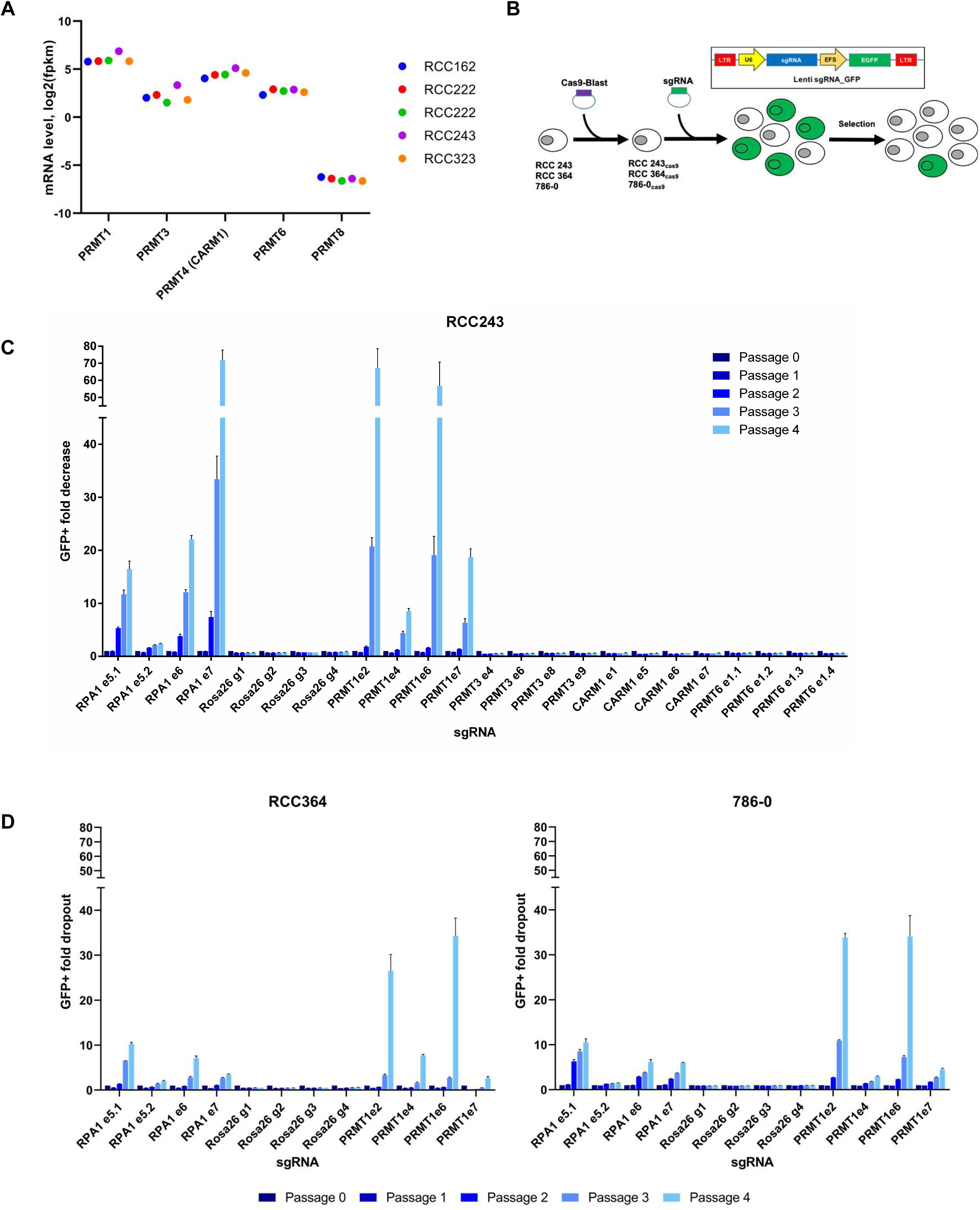
A CRISPR drop out experiment demonstrates that PRMT1 is the critical dependency among type I PRMTs in ccRCC. **a,** Type I PRMT mRNA expression in ccRCC patient-derived cell lines (each dot represents an individual cell line); fpkm, fragments per kilobase of transcript per million mapped reads. **b,** Schematic representation of the CRISPR-Cas9 competition assay used to determine the functional importance of each type I PRMT enzyme for proliferation and viability in ccRCC cells. **c,** Mean fold-change values (± SEM, n=3) in the percentage of GFP+ cells, relative to the percentage of GFP+ cells at passage 0 in cell line RCC243 for indicated type I PRMT sgRNAs. **d,** Mean fold-change values (± SEM., n=3) in the percentage of GFP+ cells, relative to the percentage of GFP+ cells at passage 0 in cell lines RCC364 and 786-0 for PRMT1 and control sgRNAs.

Briefly, cell line RCC243 was engineered to constitutively express a functional, humanized Cas-9 nuclease (RCC243_Cas9_, Supplemental Figure 3) and then subsequently transduced with GFP-expressing guide RNA (gRNA) vectors at a low multiplicity of infection (MOI). This created a mixed population of GFP+/gRNA+ and GFP-/gRNA-cells. Four gRNA’s were selected from the Broad Institute’s ‘Brunello Library’^33, 34^ for each of *PRMT1*, *PRMT3*, *PRMT4* (*CARM1*) and *PRMT6* (Supplementary Table 1). *PRMT8* was excluded due to its negligible expression levels in our ccRCC cell lines. Four additional guides were included that target each of a negative control (the human ROSA26 locus – a genetic safe harbour^35^) and a positive control (*RPA3* - an essential replication protein). Following gRNA transduction and for four subsequent passages (cells passaged at a 1:4 ratio every 4 days), the GFP+ population was monitored *via* analytical flow cytometry and the ratio of GFP+ to GFP-cells was calculated. As expected, cells with gRNAs targeting the humanized ROSA26 locus persisted in culture while cells with gRNAs targeting *RPA3* were depleted by magnitudes of 2.4 to 81.7-fold (Figure 2C). Conversely, over the course of four passages, the fold depletion for cells harbouring gRNAs that target *PRMT3*, *PRMT4* (*CARM1*) and *PRMT6* were negligible and consistent with the negative control guides. However, RCC243_cas9_ cells transduced with gRNAs targeting *PRMT1* were noticeably depleted in culture with dropout rates ranging from 7.7 to 81.9-fold. These results demonstrate that repeated mutations in *PRMT1* induced by Cas9 nuclease led to decreased cell fitness and a resulting drop-out phenotype from the overall population, while mutations in other type I PRMTs were tolerated and did not impact proliferation or viability.

To extend these results, we engineered two additional cell lines to constitutively express Cas-9 nuclease (RCC364_Cas9_ and 786-0_Cas9_, Supplemental Figure 3) and repeated the experiment with gRNA’s directed against *PRMT1* and the relevant negative and positive controls (Figure 2D). As with RCC243_cas9_, gRNA directed mutagenesis of *PRMT1* resulted in reduced cell fitness and GFP+/GFP-fold decreases comparable to those of the positive control protein *RPA3* in both cell lines. We thus conclude that PRMT1 is the critical dependency among type I PRMTs in ccRCC.

### Knockdown of PRMT1 in ccRCC phenocopies MS023 treatment and overexpression of two major PRMT1 isoforms results in drug resistant phenotypes

To supplement our CRISPR-Cas9 results, we also engineered two ccRCC cell line models to express doxycycline-inducible short hairpin RNAs (shRNAs) targeting either *PRMT1* (PRMT1-shRNA) or a ‘non-targeting’ control directed against luciferase (NT-shRNA). Upon addition of doxycycline, we observed strong depletion of PRMT1 at day 3 and day 6 in both RCC243_PRMT1-shRNA_ and 786-0_PRMT1-shRNA_ (Figure 3A). As with MS023 treatment, we also noted a marked decreased in the levels of general aDMA species in PRMT1-depleted cells relative to NT-shRNA controls, no-doxycycline treated controls and parental non-transduced cell lines (Figure 3A). To assess the impact of PRMT1 specific depletion on the growth of each cell line, proliferation was measured over a period of 10 days using the Incucyte® Live-Cell Analysis System. PRMT1 depletion significantly inhibited *in vitro* cell growth relative to NT-shRNA and non-induced controls (Figure 3B).

**Figure 3:**
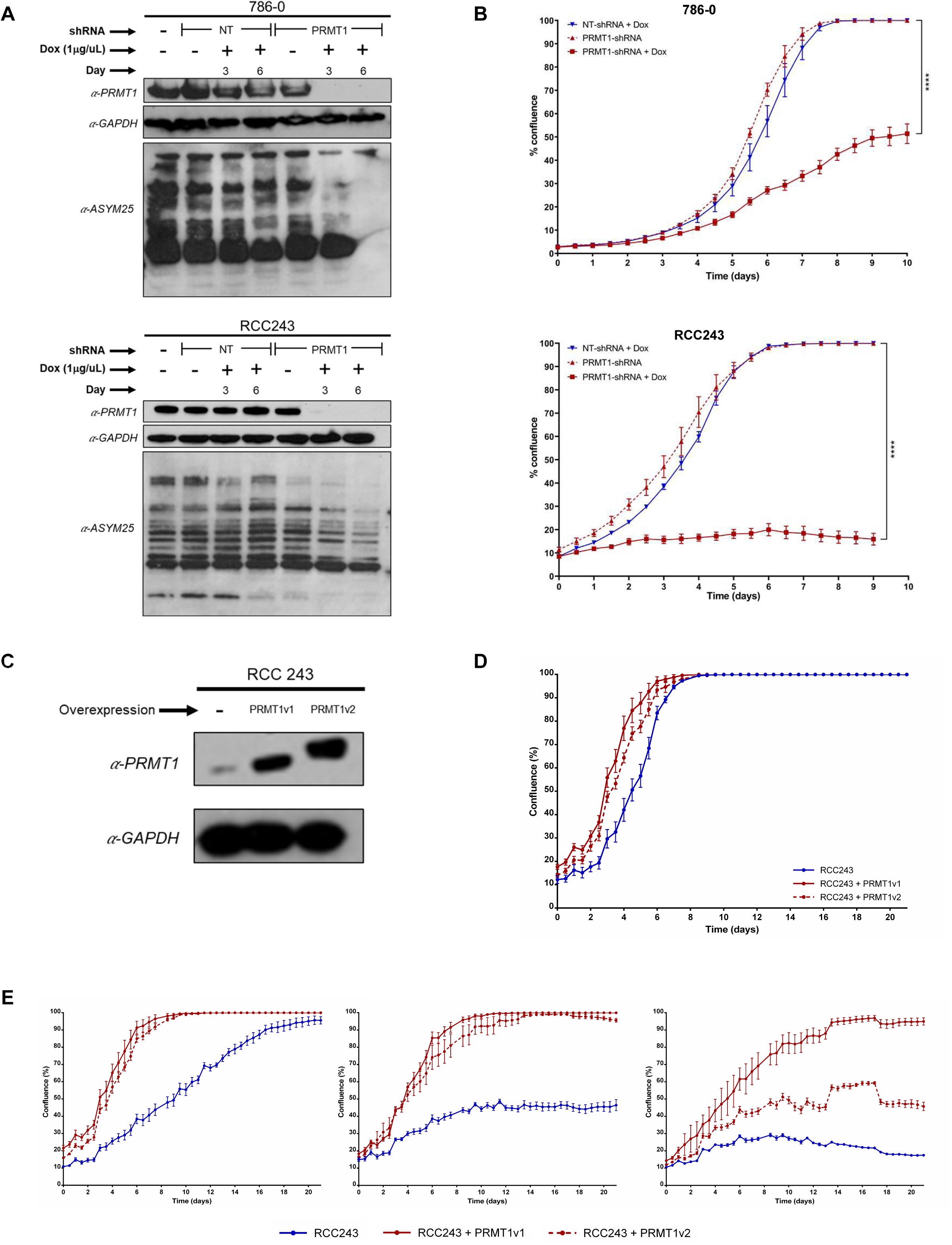
PRMT1 knockdown phenocopies MS023 treatment and overexpression results in MS023 resistance. **a,** Western blot analysis of PRMT1 expression (top) and asymmetric di-methylarginine (ADMA) changes using the ASYM25 antibody (bottom) in 786-0 and RCC 243 cells. Cell lines were engineered to express doxycycline (Dox)-inducible PRMT1 targeting or non-targeting (NT) shRNAs and treated with or without 1.0 μg/mL Dox for 3 and 6 days. **b,** Cell line growth curves (confluence measured in Incucyte® Live-Cell Analysis System) of 786-0 and RCC243 cells expressing PRMT1 targeting or NT shRNAs, with or without Dox. Data are presented as the mean ± SEM and p-values are calculated by 2-way ANOVA with repeated measures and Sidak’s multiple comparisons test. **c,** Western blot analysis of RCC243 cell line engineered to overexpress PRMT1 isoforms PRMT1v1 (ENST00000391851.8) and PRMT1v2 (ENST00000454376.7). **d,** Cell line growth curve of RCC243 cells engineered to overexpress PRMT1 isoforms. **e,** Cell line growth curves of RCC243 cells engineered to overexpress PRMT1 isoforms in the presence of 2.5 μM, 5 μM and 10 μM MS023. Data are presented as the mean ± SEM.

To further validate ccRCC’s dependence on PRMT1, we engineered patient-derived cell line RCC243 to constitutively overexpress two predominant isoforms of the PRMT1 enzyme: PRMT1v1 (ENST00000391851.8) and PRMT1v2 (ENST00000454376.7). After verifying overexpression *via* immunoblotting (Figure 3C) and confirming that said over expressions do not impact the growth dynamics of transduced cell lines relative to parental counterparts (Figure 3D), we exposed each cell line to various doses of MS023. Over expression of PRMT1v1 and, to a lesser extent, PRMT1v2 restored the aggressive proliferative phenotype seen in parental lines even in the presence of high MS023 doses up to 10µM (Figure 3E). Taken together, these data indicate that PRMT1 is a novel, targetable, therapeutic vulnerability in ccRCC.

### MS023 treatment results in a down regulation of cell cycle and DNA damage repair (DDR) pathways

To better understand the observed cellular response to MS023 treatment, our next step was to profile transcriptomic changes *via* RNA-seq analysis in cell lines RCC243 and 786-0. Following three days of treatment with 5µM MS023, each cell line was analyzed in duplicate relative to controls and a differential gene expression analysis was performed on the aggregate of both cell line replicates to look for commonly enriched molecular pathways (Figure 4A). Intriguingly, even though a total of 901 genes were found to be significantly upregulated (FDR ≤ 0.01, log_2_(FC) ≥ 2) after three days of treatment, no significant Gene Ontology (GO) biological process pathway enrichments were detected in this list. Conversely, 664 genes were found to be significantly down regulated by day 3 and a total of 33 GO biological process pathways were found with a fold enrichment of at least 2.0 and a FDR of ≤ 0.05 (Figure 4B). Prominent among these was a consistent down regulation of pathways related to the cell cycle, particularly genes involved in mitotic progression (24/33 pathways) (Figure 4B). Enrichments greater than 3-fold were noted for mitotic specific GO annotations including centromere complex assembly (GO: 0034508), mitotic cytokinesis (GO: 0000281), mitotic spindle organization (GO:0007052), mitotic sister chromatid segregation (GO: 00000070) and mitotic nuclear division (GO:0000278). Specific transcripts in these enrichments include those encoding the mitotic checkpoint protein BUB1B and a suite of centromere proteins (CENP) that play a critical role in kinetochore formation, mitotic progression, and chromosome segregation, including CENPA and CENPI (Figure 4C).

**Figure 4:**
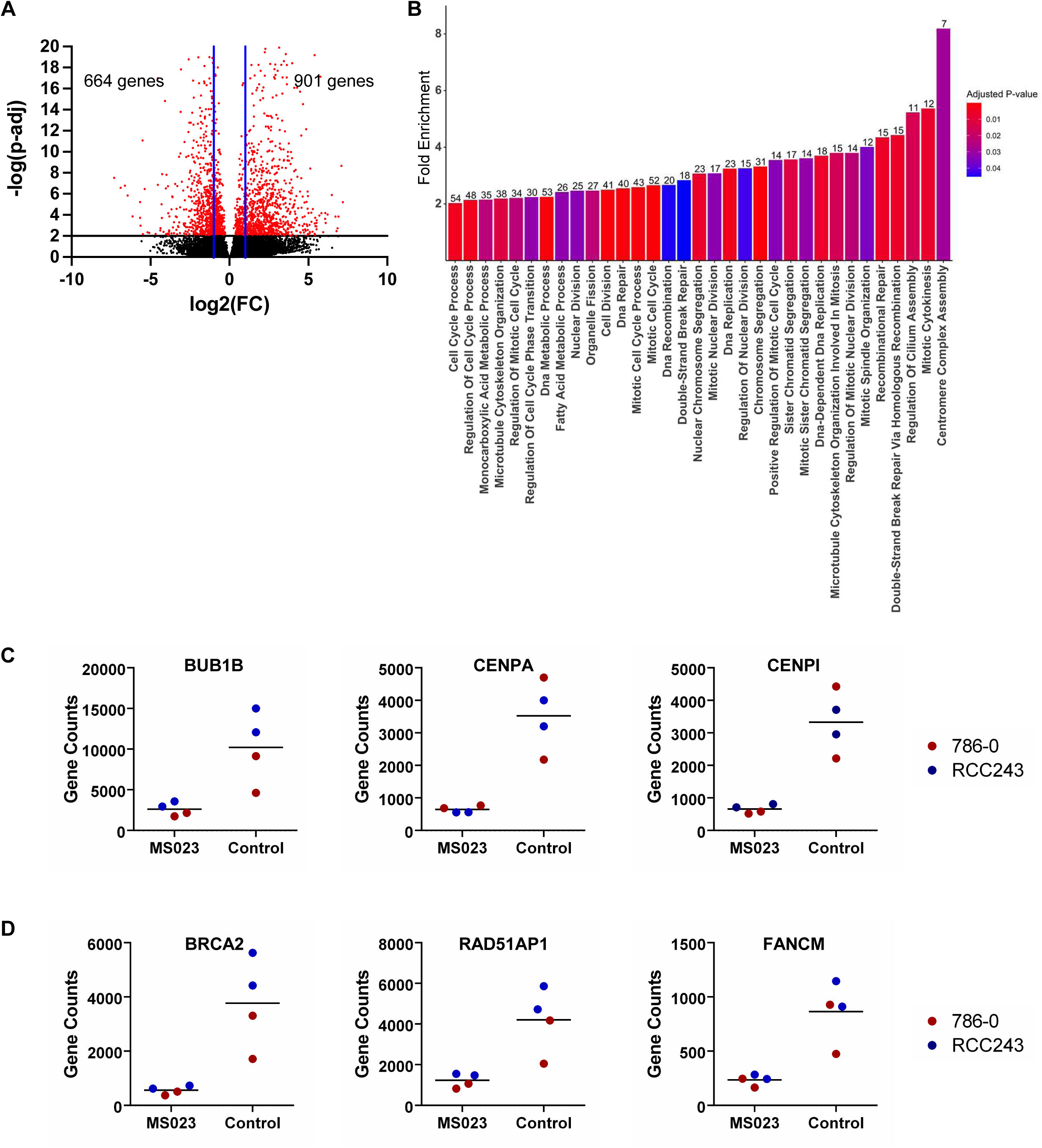
MS023 treatment leads to down-regulation of genes associated with cell cycle and DDR pathways. **a,** Volcano plot of log_2_fold-change for genes significantly downregulated (red, left) or upregulated (red, right) following 3 days of 5 μM MS023 treatment. **b,** Enrichment analysis for gene ontology (GO) biological processes on significantly down regulated genes after 3 days of MS023 treatment. Analysis conducted using Fisher’s exact test; number of genes in down regulated list per GO biological process listed above each respective bar. **c,** Gene counts for mitotic checkpoint protein BUB1B and centromere proteins CENPA and CENPI in RCC243 and 786-0 cells following 5μM MS023 treatment for 3 days. **d,** Gene counts for DNA damage proteins BRCA2, RAD51AP and FANCM in RCC243 and 786-0 cells following 5μM MS023 treatment for 3 days.

Also prominent among the enriched pathways were those related to DNA damage repair (DDR). GO annotations for DSB repair *via* homologous recombination (GO:0000724), and recombinational repair (GO:0000725) were detected at enrichment levels greater than 4-fold (Figure 4B). Included among the proteins in these lists were important players in homologous recombination (HR) including the breast cancer type 2 susceptibility protein (BRCA2) and the RAD51-associated protein (RAD51AP1). Additionally, the Fanconi anemia group M protein (FANCM), a key player in the regulation of DNA interstrand cross-link (ICL) repair was also significantly downregulated (Figure 4D).

### Type I PRMT inhibition results in a stalled cell cycle, decreased expression of DDR proteins and an accumulation of DSBs

To further assess the apparent transcriptomic down regulation of cell cycle related pathways, immunoblotting for specific mitotic proteins was performed after treatment with various doses of MS023 for 3 days. Additionally, we assessed the relative cell population actively undergoing division in a time course experiment using an anti-phospho-Histone H3 (Ser10) antibody (Anti-H3S10p), a well characterized marker of mitosis^36^. An MS023 dose dependent reduction in levels of BUB1B, CENPA and CENPI was detected after three days of treatment (Figure 5A) and immunoblotting with Anti-H3S10p in RCC243 following 3, 5 and 7 days of 5 µM MS023 exposure confirmed a decline in the relative cell population actively undergoing mitosis relative to controls (Figure 5B). However, a DNA content analysis of RCC243 cells over a treatment period of 9 days revealed no accumulation in any specific phase of the cell cycle with prolonged exposure to 5µM MS023 (Figure 5C). After 5 days of treatment, a population of cells with sub-2N DNA content (i.e. cells with DNA fragmentation) began to appear, while the number of cells in other phases of the cell cycle began to lag control conditions. By day 9, the sub-2N population became dominant with few cells remaining in other cell cycle phases (Figure 5C). Overall, MS023 treatment appears to induce a cytostatic phenotype that precedes cell death.

**Figure 5:**
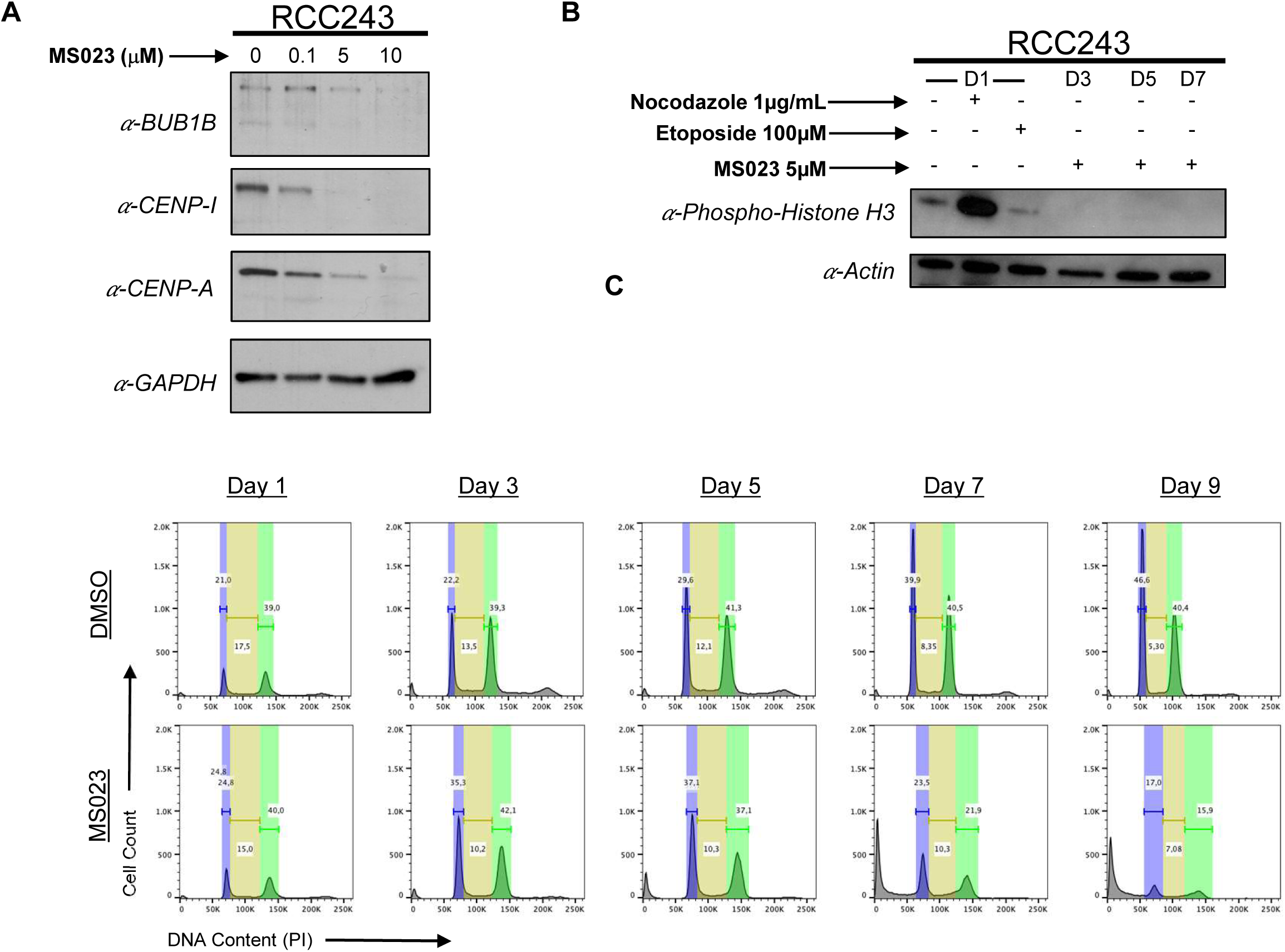
MS023 treatment results in a stalled cell cycle and eventual cell death. **a,** Western blot analysis of mitotic checkpoint protein BUB1B and centromere proteins CENPA and CENPI in RCC243 cells treated with 0 μM, 0.1 μM, 5 μM, and 10 μM of MS023 for 3 days. **b,** Western blot analysis of phospho-histone H3 in RCC243 cells treated with 0.1 μg/mL Nocodazole (beta-tubulin disruptor and mitotic staller), 100 μM etoposide (DNA damaging agent) and 5 μM MS023 for 1, 3, 5 and 7 days. **c,** Flow cytometric histograms showing cell cycle progression of RCC243 cells in response to 5 μM MS023 inhibition over 9 days.

Similarly, to assess the impact of MS023 treatment on DDR proteins, we performed specific immunoblotting in RCC243 for the down regulated targets identified in our transcriptomic analysis BRCA2, RAD51AP1 and FANCM (Figure 6A). We expanded the repertoire of immunoblotting targets to include other key players in the Homologous Recombination and Fanconi Anemia DDR pathways (Figure 6A). For all targets assessed, a dose dependent reduction in protein levels was noted with 3 days of MS023 treatment indicating a possible compromise in the integrity of these critical repair pathways.

**Figure 6:**
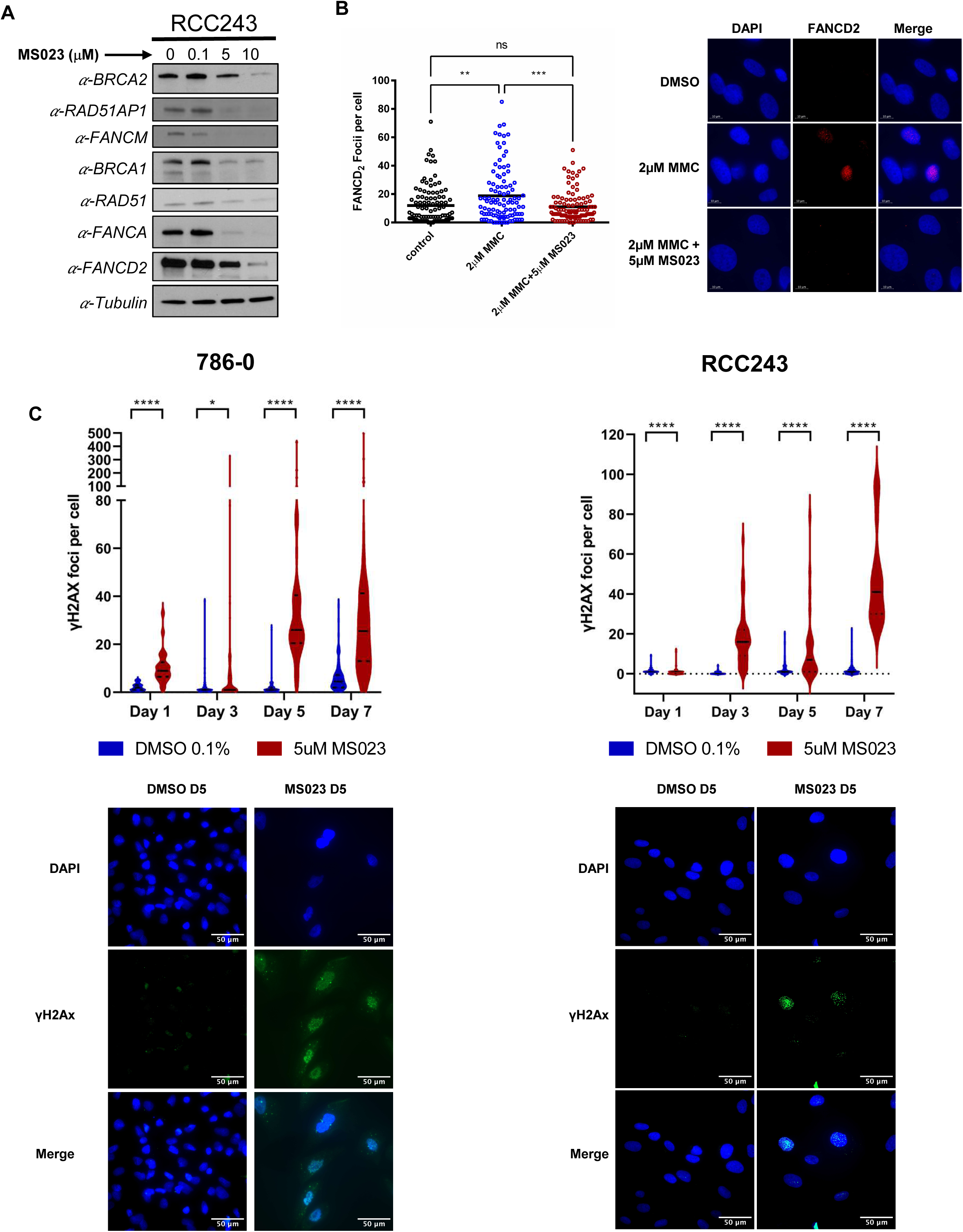
MS023 treatment decreases DDR proteins, impedes the formation of FANCD2 foci upon mitomycin C treatment, and causes accumulation of DSBs. **a,** Western blot analysis of indicated of indicated proteins in RCC243 cells treated with 0 μM, 0.1 μM, 5 μM and 10 μM of MS023 for 3 days. **b,** Scatter plots of FANCD2 foci in RCC243 cells treated with and without 5 μM MS023 for 3 days and then exposed to 2μM Mitomycin for 24 hours to induce interstrand cross linking. P-values are calculated by one way ANOVA with repeated measures and Sidak’s multiple comparisons test. Representative images are shown beside the plot. ns, not significant; **, p<0.01; ****, p<0.001. n = 103 to 107 cells analyzed. **c,** Violin plots of γH2AX foci in RCC243 and 786-0 cells treated with and without 5 μM MS023 for indicated times. P-values are calculated by unpaired t-test. Representative images from Day 5 of treatment are shown below each plot. ns, not significant; *, p<0.05; **, p<0.01; ****, p<0.001. n = 16 to 44 cells analyzed.

To investigate the integrity of DNA ICL repair, we performed immunofluorescent staining for FANCD2 foci, a hallmark of a functional FA pathway. RCC243 cells were treated for 3 days with 5µM MS023 and then exposed to the interstrand cross link agent mitomycin C (MMC). Consistent with the observed reduction in levels of FA proteins, treatment with MS023 significantly reduced the number of FANCD2 foci following exposure to MMC relative to controls, indicating a compromised ability to handle such damage (Figure 6B).

To further evaluate the potential for a DDR compromised phenotype in cell lines RCC243 and 786-0, we performed immunofluorescent foci staining for γH2AX, a marker of DSBs. A significant increase in the number of γH2AX foci with 5µM MS023 treatment was noted over time relative to control conditions indicating an accumulation of unresolved DSBs (Figure 6C). We also analyzed DNA repair efficiency following ionizing radiation (IR) exposure using the comet assay. This assay was performed in cell line RCC243 under neutral conditions to specifically quantify the removal of DSBs. Relative to controls, we observed that cells treated with MS023 repaired DNA breaks more slowly, signalling a defect in the cells’ DSB repair mechanisms (Supplemental Figure 4). This deficit was comparable to that seen with the use of a DNA-dependent protein kinase (DNAPK) inhibitor, a well characterized disruptor of DSB repair. Taken together our data suggests that the type I PRMTs coordinate the expression of key DDR genes and that inhibition of these enzymes leads to impaired DDR pathways, accumulation of DSBs, cell cycle arrest and eventual cell death.

### Bio-ID reveals RNA-binding proteins (RBPs) as the predominant PRMT1 interactors

To elucidate the molecular mechanisms driving PRMT1-mediated inhibition of ccRCC growth, we employed a novel, proximity-labelling (PL) methodology using miniTurbo^37^, an adaptation of the BioID technique^38^. The advantage of miniTurbo relative to the legacy BioID system is the relative speed at which the PL reaction takes place^37^. Although PRMT1 interactome data has previously been published^22, 23, 39, 40^, these studies have relied on standard affinity purification of PRMT1 interacting partners coupled with mass spectrometry (AP-MS), or analysis of differentially methylated proteins *via* AP-MS or isotope labelling techniques such as methyl-SILAC. The establishment of differentially methylated states for substrate interactors of PRMT1 has traditionally been achieved through chemical inhibition of type I PRMTs, which lacks specificity for PRMT1, or *via* genetic depletion of the enzyme, which is prone to off-target effects. Standard AP-MS techniques also yield a number false negatives for low abundance proteins and for interactors that are difficult to solubilize, such as chromatin associated proteins, nuclear matrix and other insoluble cellular compartments^38^.

For our PL method, a genetic fusion was created between the PRMT1v1 isoform and an engineered, promiscuous mutant of the Escherichia coli-derived biotin ligase BirA known as miniTurbo. This fusion was genetically cloned into a doxycycline-inducible construct that was transduced *via* lentiviral delivery into cell lines RCC243 and 786-0 (Supplemental Figure 5). Following induction and expression of the fusion protein, exogenous biotin was added to the cell culture to initiate the covalent tagging action of the miniTurbo enzyme (Supplemental Figure 5). Thus, any protein interactors of PRMT1 that are brought within a few nanometers of the fusion receive a biotin tag. Because of the irreversible nature of these biotin labels, the cells can then be lysed under harsher buffer conditions to maximize solubilization. Labelled proteins were purified by high affinity streptavidin precipitation and identified by mass spectrometry. With this technique, we were able to define the PRMT1 interactome in ccRCC cells in a specific manner under physiological conditions and over a short window of time (90 minutes with miniTurbo).

A total of 59 high confidence PRMT1 interactors (log_2_(FC) ≥2 and ≥20 total counts) were identified across both RCC243 and 786-0 (Figure 7A). The agreement between these data sets was high with a total of 41/59 interactors common to both cell lines. GO biological process and molecular function pathway enrichment analysis of the common list revealed that the majority of identified PRMT1 interactors are proteins involved in RNA metabolism and processing (Figure 7B). This is consistent with previously published reports and underscores a crucial role for PRMT1 as a regulator of RNA effector molecules involved in splicing, translation, and other RNA regulatory activities^22, 23, 41, 42^. Intriguingly, one of the top interactors in both RCC243 and 786-0 was Bcl2-associated transcription factor (Btf or BCLAF1). BCLAF1 is serine-arginine-rich (SR) RNA processing factor that has recently been implicated as a regulator of splicing and cytoplasmic shuttling for cellular DDR pathways^43, 44^. To determine the methylation status of BCLAF1, immunoprecipitation (IP) was done using a BCLAF1 specific antibody followed by aDMA specific immunoblotting (Figure 7C). We note that with 5 µM MS023 treatment for 3 days the methylation level for this PRMT1 interactor was markedly decreased, verifying BCLAF1 as a PRMT1 substrate (Figure 7C).

**Figure 7:**
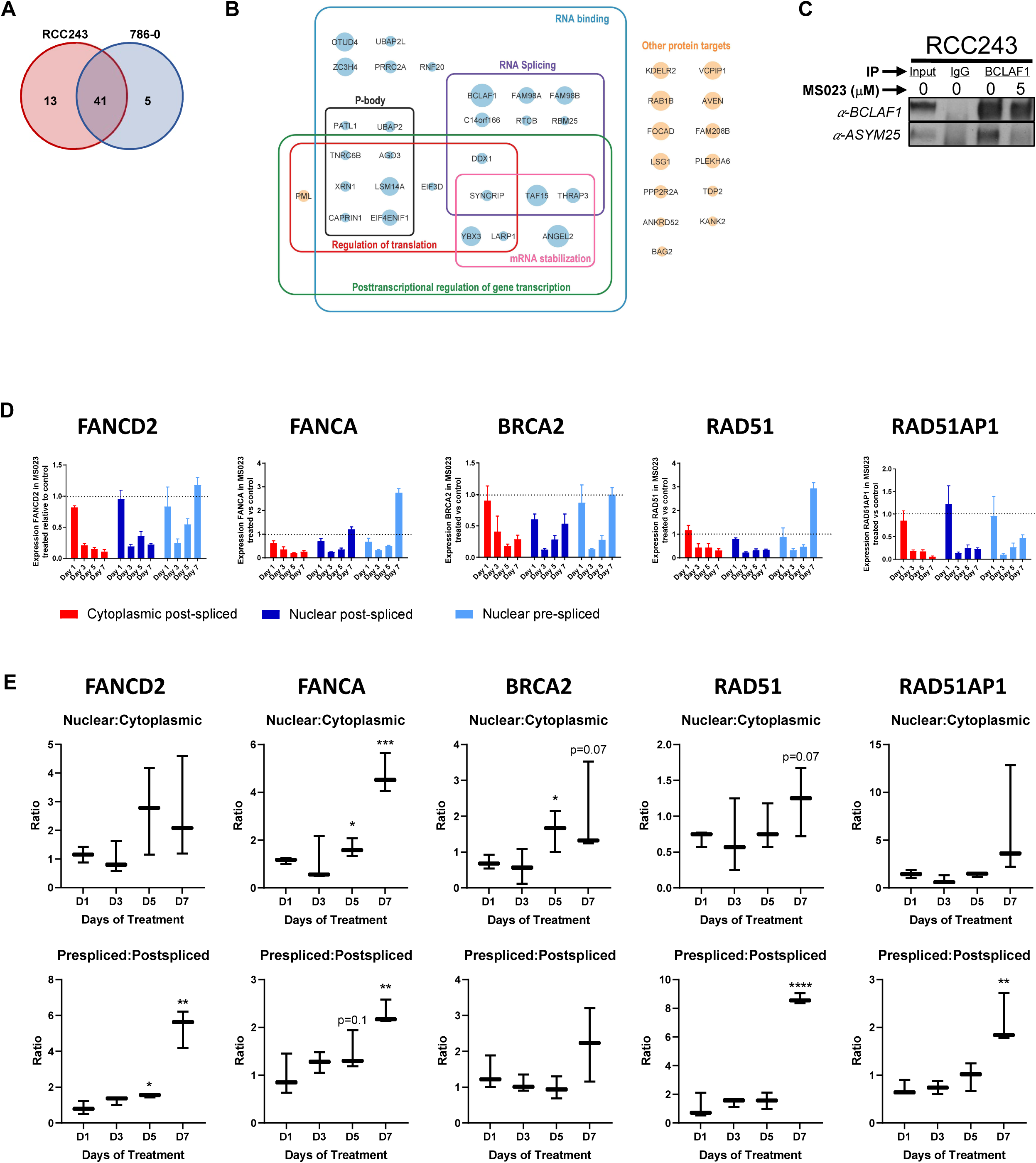
Bio-ID reveals RNA-binding proteins as predominant PRMT1 interactors, including BCLAF1 and THRAP3. **a,** Venn diagram of the 59 total high confidence PRMT1 interactors identified in mini-Turbo Bio-ID experiment (log2(FC) ≥2 and ≥20 total counts) in RCC243 and/or 786-0. **b,** Summary diagram of the 41 PRMT1 interactors identified in both cell lines. Protein names were imported into Cytoscape 3.9.1 for visual representation and enrichment analysis was carried out using the STRING Enrichment app using the categories GO biological process, GO molecular function and COMPARTMENTS. Circle size corresponds to fold-change in peptide counts between PRMT1-expressing vs control-miniTurbo cells. The majority of PRMT1 interactors correspond to RNA-binding proteins (blue circles), with the remainder corresponding to other proteins that do not fall into any significantly enriched categories (orange circles). **d,** RCC243 cells were incubated with 5 μM of MS023 or 0.1% DMSO for 3 days, lysed and anti-BCLAF immunoprecipitations were performed. The bound proteins were separated by SDS-PAGE and immunoblotted with the indicated antibodies. **e,** mRNA levels for indicated genes in denoted cellular fraction after exposure to 1, 3, 5 and 7 days treatment with 5 µM MS023. **f,** Ratio of nuclear/cytoplasmic post-spliced transcripts and ratio of nuclear pre-spliced to nuclear post-spliced transcripts. mRNA expression levels assessed via qRT-PCR on cDNA generated from DNAse treated RNA samples and normalized to Actin. Graphs represent the mean of three independent experiments ±SEM. Significant differences in the ratios were assessed using Student’s two-tailed t-test in comparison to DMSO-treated cells and are indicated by ∗p< 0.05, **p<0.01.

BCLAF1 and the thyroid hormone receptor associated protein 3 (THRAP3), also identified as a high confidence PRMT1 interactor, have been identified as core members of a DNA damage-induced BRCA1-mRNA splicing complex that promotes the pre-mRNA splicing and subsequent transcript stability of a subset of genes^43^. Savage *et al* showed that this complex is recruited to genes constitutively bound by BRCA1 at exons and exon-intron boundary regions and promotes the pre-mRNA splicing and subsequent transcript stability of a number of genes involved in DDR^43^. A subsequent study showed that BCLAF1 and THRAP3 also promote RNA splicing and export, respectively, of Fanconi Anemia (FA) and HR DDR pathway transcripts independently of DNA damage including FANCL, FANCD2, BRCA2 and RAD51^44^.

Given the role of BCLAF1 and THRAP3 in the pre-mRNA splicing and nuclear export of genes involved in DDR pathways^44^, we selected a subset of disrupted DDR genes from our previously generated RNA-seq and Western blot assays and carried out qRT-PCR assays on isolated cytoplasmic and nuclear RNA using primers spanning exon-exon and exon-intron boundaries to quantify levels of pre-spliced and post-spliced RNA in each cellular compartment (Figure 7D). Efficiency of RNA fractionation in these experiments was confirmed by assaying levels of the nuclear RNA, MALAT1 (Supplemental Figure 6). We note that the qRT-PCR confirms the overall decrease in mRNA levels, both pre-spliced and post-spliced, of all of the tested genes within 3 days of MS023 treatment. In addition, there is a general trend towards accumulation of pre-spliced transcripts in the nucleus that first decreases but then increases over time, as illustrated by increasing pre-spliced to post-spliced ratios in the nuclear fraction on days 5 and 7 (Figure 7E). The nuclear:cytoplasmic ratio of post-spliced RNA also shows an increase over time for some of the tested genes (Figure 7E). This pattern is most consistent for the *FANCD2* and *FANCA* genes, in line with our FANCD2 foci data (Figure 6B). Thus overall, this data suggests that, in addition to impacting transcription of DDR proteins in ccRCC cells, MS023 treatment also impacts nucleocytoplasmic transport and RNA splicing of these genes, possibly through regulation of the BCLAF/THRAP3 complex.

### Type I PRMT inhibition and PRMT1 genetic knock down suppresses tumor growth *in vivo*

To determine if the anti-proliferative effects of type I PRMT inhibition in ccRCC cell lines translate to antitumor activity *in vivo*, we evaluated the efficacy of MS023 treatment in ccRCC xenografts. To establish a maximum tolerated dose, intraperitoneal (IP) injections ranging from 40 to 160 mg/kg of MS023 were administered, 5 days per week over a course of 3 weeks and 80mg/kg was selected as the maximum tolerable dose (≤10% body weight loss over the treatment period). Subsequent pharmacokinetic analysis of drug distribution in serum, kidney, liver, and tumor tissues demonstrated that after three consecutive days of IP injections at 80mg/kg, residual amounts of the drug could be detected in tumor tissues up to four days later (Supplemental Figure 7). Thus, to improve tolerability parameters, a dose of 80mg/kg was selected for efficacy evaluation with an administration schedule of 3 days on/4 day off.

RCC243 and 786-0 cells were implanted subcutaneously, and after tumor establishment, mice were randomized to receive either control (vehicle solution) or 80 mg/kg of MS023 *via* IP injection in our predetermined dosing schedule. Treatment was continued until controls reached the humane endpoint. Significant inhibition of tumor growth was noted for both RCC243 and 786-0 xenografts relative to controls (Figure 8A). Global inhibition of the aDMA mark was also noted in xenograft tumor tissue as evaluated by immunoblotting in RCC243 (Figure 8B) suggesting that tumor growth inhibition coincides with type I PRMT inhibition. To extend these results, we also tested the tumor inhibition potential of MS023 in two patient derived xenograft (PDX) models (generated by direct implantation of patient-derived tumor tissues into immunocompromised mice). Similar to the cell line derived models, growth of RCC63 and RCC243 PDX tumors were significantly reduced by MS023 administration relative to controls (Figure 8C).

**Figure 8:**
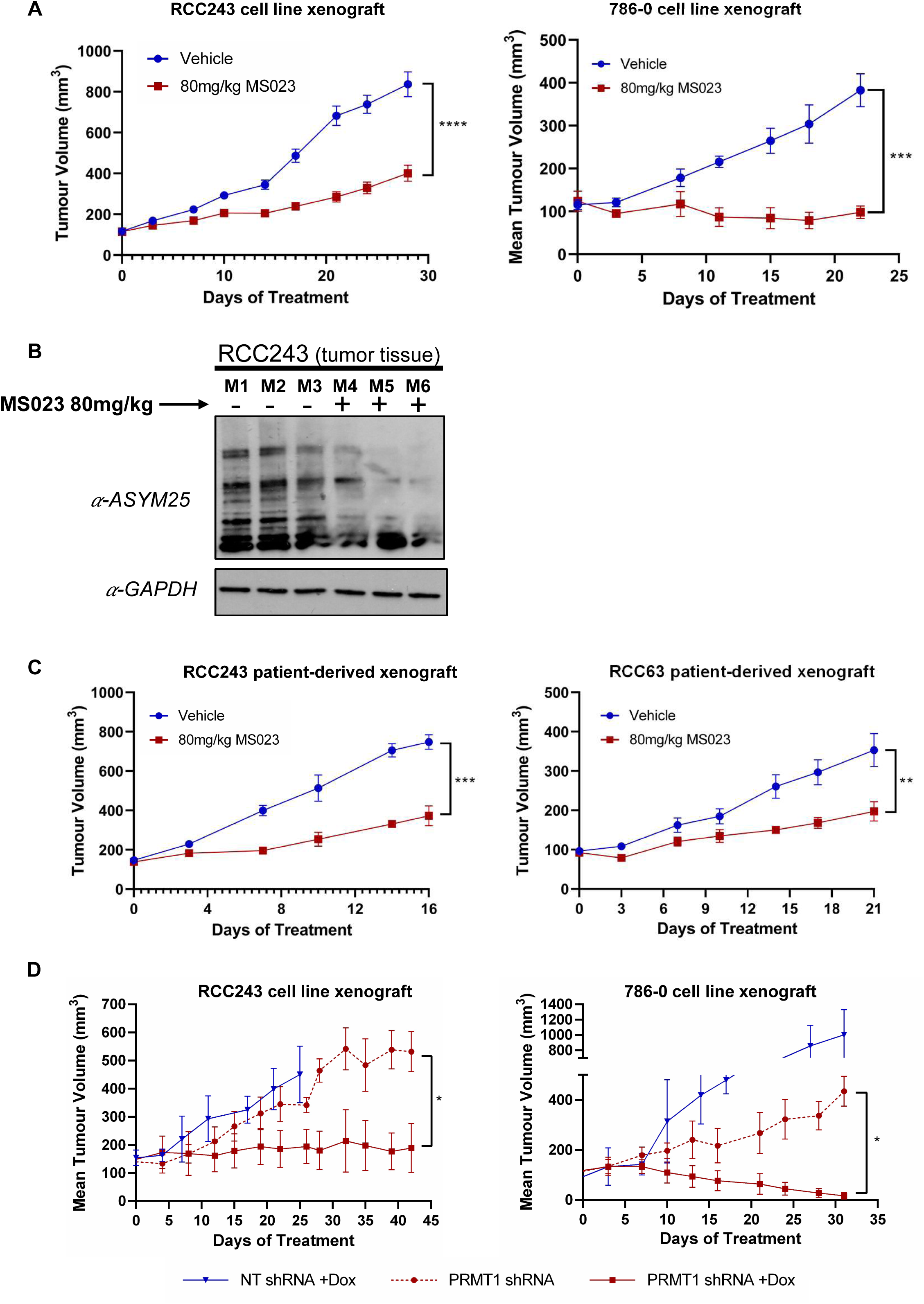
MS023 treatment and PRMT1 knockdown suppress tumor growth *in vivo*. **a,** Tumor growth curves of RCC243 and 786-0 cell line xenograft models treated with MS023 at 80 mg/kg or vehicle control, QD 3 on/4 off. Data are presented as the mean ± SEM and p-values are calculated by 2-way ANOVA with repeated measures and Sidak’s multiple comparisons test. *n* = 5 to 10 mice/group. ***, p<0.005; ****, p<0.001. **b,** Western blot analysis of ADMA in RCC243 tumor lysates on day 30. **c,** Tumor growth curve of RCC63 and RCC243 patient-derived xenograft (PDX) models treated with MS023 at 80mg/kg or vehicle control, QD 3 on/4 off. Data are presented as the mean± SEM and p-values are calculated by 2-way ANOVA with repeated measures and Sidak’s multiple comparisons test. *n* =5 to 10 mice/group. **, p<0.01; ****, p<0.001. **d,** Tumor growth curves of RCC243 and 786-0 cell line xenografts expressing Dox-inducible *PRMT1*-targeting or NT control shRNAs (*n* = 3 to 5 mice/group). Mice were randomized to either Dox supplemented water (1mg/mL dox in water) or normal water upon tumor establishment (100-200 mm^3^). Data are presented as the mean ± SEM and p-values are calculated by 2-way ANOVA with repeated measures and Sidak’s multiple comparisons test. *, p<0.05.

Finally, to further verify that PRMT1 is the specific type I enzyme responsible for tumor growth inhibition, we evaluated the effect of PRMT1 genetic depletion on tumor growth *in vivo*. RCC243_PRMT1-shRNA_/RCC243_NT-shRNA_ and 786-0_PRMT1-shRNA_/786-0_NT-shRNA_ cells were implanted subcutaneously, and tumors were established. Mice were then randomized to receive a normal diet, or one supplemented with doxycycline to induce genetic depletion. Like MS023 treatment, depletion of PRMT1 in mice bearing tumors from both cell lines resulted in significantly inhibited tumor growth relative to NT-shRNA and non-induced controls (Figure 7D). These results from preclinical models suggests that PRMT1 inhibition holds promise as a potential therapeutic strategy in the treatment of ccRCC.

## Discussion

A defining feature of ccRCC tumors is their stubborn resistance to radiation and traditional chemotherapeutics, a hallmark that has left few treatment options beyond surgical resection. Detailed genomic analyses of this cancer have revealed a complex and heterogenous molecular picture, underpinned by two common themes: a near universal dependence on distorted hypoxia signalling and distorted chromatin regulation as key drivers of tumorigenesis. In this study, we sought to exploit the latter of these dependencies using a targeted, epigenetic focused chemical screening approach in well characterized patient-derived models of ccRCC. Through this unbiased methodology, we identified the type I PRMT inhibitor MS023 as an attenuator of cell growth. Using orthogonal genetic techniques, we further identified PRMT1 as the specific dependency responsible for growth arrest among the type I enzymes. We show that MS023 administration and PRMT1 inhibition result in widespread loss of aDMA species including the histone mark H4R3me2a. Our transcriptomic data demonstrates pronounced downregulation of key mitotic and DDR proteins that coincide with marked inhibition of the cell cycle, compromised DDR pathways, and an accumulation of DSBs. These observations are consistent with previous reports that also show cell cycle defects and DNA damage resulting from PRMT1 deficiencies. Encouragingly, our preclinical *in vivo* models also demonstrate significant tumor growth inhibition following MS023 administration and PRMT1 genetic depletion, suggesting this enzyme represents a viable therapeutic target for potential clinical development.

To further delineate relevant molecular pathways at play, we employed a novel PL proteomics technique to describe the interacting partners of PRMT1 in ccRCC cells. Consistent with previous reports, the bulk of interactors identified were RNA binding proteins (RBPs) that regulate key RNA metabolic activities including mRNA transcription, splicing, transport, translation, and turnover. Of the top PRMT1 interactors described, BCLAF1, is known to directly influence the splicing and transport of important DDR protein transcripts in conjunction with another top interactor, THRAP3^43, 44^. These splicing targets include key members of the Homologous Recombination and Fanconi Anemia DDR pathways^43, 44^. Strikingly, many of these same proteins were also downregulated in response to type I PRMT inhibition. Moreover, our splicing analysis demonstrated a down regulation of nucleocytoplasmic transport and RNA splicing following prolonged exposure to MS023 for the DDR genes identified in our study. While we were able to confirm differential asymmetric di-methylation of BCLAF1 in the presence of MS023, the exact influence of this PTM on the ability of BCLAF1 and THRAP3 to act as splice regulators is unknown. Methylation of other RBPs has been shown to mediate important mechanisms like nucleocytoplasmic shuttling, protein-protein interactions, and nucleic acid binding^18, 45^. Conceivably, PRMT1 inhibition and the resulting loss of aDMA PTMs on BCLAF1 may impact its ability to effectively regulate splicing, resulting in compromised transcript quality and protein disruptions of the DDR targets described in this study.

Although a collapse in splicing is one plausible explanation for the DNA damaged/growth arrested phenotypes seen with MS023 treatment, it should be noted that PRMT1 also targets an array of specific DNA damage proteins, including BRCA1, hnRNPK, hnRNPUL1, MRE11 and 53BP1^46–49^. These proteins are key players in DSB repair and their PRMT1-mediated aDMA PTMs are known to affect their abilities in this regard. MS023 treatment could plausibly lead to unresolved DNA damage by impairing the function of these proteins leading to cell cycle arrest and eventual cell death. However, we note that key proteins from the Fanconi Anemia pathway, a related yet mechanistically separate DDR pathway specialized for the removal of DNA interstrand crosslinks, are also critically downregulated by MS023 treatment. The functional compromises we described in this pathway are not readily explained by changes in aDMA PTMs for MRE11, 53BP1, hnRNPK, hnRNPUL1 or BRCA1.

A relative black box in our mechanistic exploration of PRMT1 in ccRCC revolves around the influence of its histone substrate, H4R3me2a on the overall cellular transcriptome and resulting proteome. Despite recent advancements in high throughput sequencing technologies, the epigenomic profiling of arginine histone substrates has remained an outstanding challenge owing in part to a dearth of high quality, validated antibodies against methylated arginine histones^50^. We know that H4R3me2a is associated with transcriptional activation, but to our knowledge no successful genome wide mapping of this PRMT1 mediated mark has yet been accomplished. Since our data suggests that MS023 treatment diminishes this histone mark in our disease models, we must consider the possibility that H4R3me2a-mediated transcriptional changes also underly the observed growth arrested phenotypes.

Finally, we note that type I PRMT inhibition resulted in varying levels of growth inhibition across our ccRCC cell line panel, a result that is consistent with previously reported type I PRMT efficacy studies in other cancer types^51^. This spectrum of sensitivity likely results from the heterogenous genomic and epigenomic character of our cell lines. Previous reports have linked PRMT1 sensitivity to factors like MTAP loss, SRSF mutations and gene expression signatures enriched for interferon response and antiviral signaling^21, 23, 51^. While these genetic factors are uncommon in ccRCC, the conserved evolutionary driver subtypes seen clinically most likely play a role in the cells’ sensitivity to PRMT1 inhibition. For example, BAP1 driven ccRCCs are known to have inherently compromised BRCA-mediated DDR pathways and may exhibit enhanced sensitivity to further DDR perturbations, while SETD2 driver mutations already display splicing defects and may have enhanced responses^11^. Further investigation of PRMT1 sensitivity in ccRCC subtypes is warranted and would help facilitate a potential molecular-based stratification for drug responses.

To summarize, our findings reveal the central role of PRMT1 as a regulator of RNA metabolism, including splicing, and add to the growing body of knowledge linking splicing disruptions to the DNA damage response. We demonstrate that PRMT1 plays a critical role in the maintenance and growth of ccRCC and may represent a key therapeutic vulnerability. The potential also exists for this target to synergize with current DNA-damaging therapies including radiation and traditional chemotherapeutics. In addition, the inability of MS023 treated cells to repair DSBs suggests the possibility of synergy with other DNA repair targeted drugs such as PARP inhibitors. Finally, the induction of DNA damage may lead to increased neoantigen load and/or stimulate the cGAS/STING pathway^52^, leading to increased sensitivity to immune checkpoint blockade. Further studies are needed to investigate these possibilities.

## Materials and Methods

### Cell Lines & Cell Culture

The human kidney cancer cell line 786-0 was obtained from American Type Culture Collection (ATCC). Cell lines RCC22, RCC162, RCC222, RCC243, RCC323, RCC364 and RCC407 were generated as previously described in Lobo et al^14^. The continued use of these patient-derived lines is approved under UHN Research Ethics Board approval, protocol #15-9559. All cell lines in this study were routinely cultured in Iscove’s Modified Dulbecco’s Medium (IMDM, Wisent #319-105-CL) supplemented with 10% FBS (Thermo #12483020), and 1% penicillin/streptomycin (Wisent #450-201-EL) in a humidified incubator at 37°C with 5% CO_2_, 2%O_2_. Cells were plated in culture flasks coated with rat tail collagen type I (5ug/cm^2^; Thermo #A1048301) and passaged no more than 20 times. Cell cultures were monitored for mycoplasma infection (Universal Mycoplasma Detection Kit, Cedarlane #30-1012K) and cell culture identity verification was done by short tandem repeat profiling (GenePrint® 10 System, Promega #B9510).

### Targeted DNA sequencing

DNA was extracted using a QiaAMP DNA mini kit (Qiagen). Libraries were constructed using a KAPA Hyper Prep kit using custom unique molecular identifier BIOO Scientific NextFlex adapters to barcode the samples, as instructed by the manufacturer. Target capture for the genes of interest was carried out using custom xGen® Lockdown® probes purchased from IDT. Following end repair, A-tailing and adapter ligation, Agencourt AMPure XP beads were used for library clean up and ligated fragments were amplified 4 cycles using 0.5 μM custom unique xGen Predesigned Hybrid Panel index primers (IDT). Post amplification cleanup was performed using Agencourt AMPure XP beads. Final library quality control was performed using a Bioanalyzer 2100. Sequencing was done on an Illumina HiSeq2000 using a 100-cycle paired-end protocol.

### Epigenetic chemical probe screen

All ‘epiprobe’ compounds were purchased from Cayman Chemical, Millipore-Sigma or MedChemExpress. The detailed resources for each compound are listed in supplementary table 1 (Table S1) the additional file. Chemical purity was validated at the SGC at more than 99%. Cell lines were plated at the density of 2500 cells/well (except 786-0 which was plated at 150 cells/well) in 96-well plates and allowed to adhere. Epiprobes were dissolved in DMSO and added to achieve a final concentration as indicated in Figure 1B. Each plate contained three replicates per compound and three replicates of a 0.1% DMSO control condition. After seven days exposure to each probe, the cell permeable, far-red DNA fluorescent dye DRAQ5^TM^ was added to the wells and fluorescence was quantified using the LI-COR imaging system. Data was normalized to the DMSO control wells and the average log_2_(probe/DMSO) readings are presented.

### Western Blot Analysis

Cells were lysed and proteins were extracted in ice cold RIPA buffer (150mM NaCl, 1.0%NP40, 0.5% deoxycholate, 0.1% SDS, 50mM Tris pH 8.0) supplemented with HaltTM Protease Inhibitor Cocktail (Thermo # 78425). Protein concentrations were determined using the PierceTM BCA Protein Assay Kit according to the manufacturer’s instructions. Protein concentrations were normalized to 15 μg each, and samples were denatured in Laemmli sample buffer (BioRad #1610737) supplemented with 5% β-mercaptoethanol and boiled at 95°C for 5 min. Proteins were separated on 4–15% Mini-PROTEAN® TGXTM Precast Protein Gels (BioRad, #4568084) using the Mini-PROTEAN® Electrophoresis System (BioRad). Separated proteins were then transferred to Immobilon®-P PVDF Membranes (Milipore Sigma #ISEQ00010) using the BioRad Transblot SD Semi-dry Transfer Cell as per manufacturer’s recommendations.

Membranes were blocked in 5% milk Tris-buffered saline with 0.1% Tween® 20 Detergent (TBST) and incubated overnight with primary antibody. Antibodies used: Asymmetric Di-Methyl Arginine ASYM25 (Sigma, #09-814, 1:1000), Tubulin (Santa Cruz #sc-5286, 1:1000), Histone H4 (Abcam, #ab174628, 1:1000), Histone H4R3me2a (Active Motif, #39705, 1:1000), PRMT1 (Thermo, PA5-17299, 1:1000), GAPDH (Santa Cruz, #sc-69778, 1:1000), BUB1B (Cell Signalling Technology (CST), #4116S, 1:1000), CENP-I (CST, #49426S, 1:1000), CENP-A (CST, #2186T, 1:1000), Phospho-Histone H3 (Ser10) (Millipore, #06-570, 1:1000), Actin (Abcam #ab7817, 1:1000), BRCA2 (CST, #107415, 1:1000), RAD51AP1 (Thermo/Proteintech, #11255-1-AP, 1:1000), FANCM (Bethyl Labs, #A302-637A, 1:1000), BRCA1 (CST, #14823S, 1:1000), RAD51 (CST, #8875, 1:1000), FANCA (CST, #14657, 1:1000), FANCD2 (CST, #16323, 1:1000), BCLAF1 (Thermo/Bethyl Labs, #A300-608A, 1:1000), CAS9 (Abcam, #ab191468, 1:1000), FLAG (Sigma, F3165, 1:1000), Streptavidin-HRP (BD Pharmingen, #554066, 1:1000).

Membranes were washed 5x in TBST and probed with species specific HRP-conjugated secondary antibody (anti-rabbit-HRP (CST, #7074S, 1:2500), or anti-mouse-HRP (CST, #7076S, 1:1000)) for 1 hour. Membranes were then rewashed 5x in TBST and developed with the chemiluminescence AmershamTM ECLTM start Western blotting detection reagent and visualized on Hyblot CL film (Cedar Lane, #DV-E3012).

### Western Blotting of tumor tissue

50 mg fragments of snap frozen tissue were homogenized using a Qia-tissue ruptor at medium speed for 30 seconds or until completely homogenous, on ice in RIPA buffer containing protease and phosphatase inhibitors. Samples were sonicated twice for 10 seconds at 50% amplitude.

### Plasmids, Transfections, Transductions and Engineered Cell Line Development

All lentiviral preparations were made via co-transfection of target and packaging plasmids in HEK293T cells using the XtremeGENE HP DNA transfection reagent (Sigma, #6366236001). Packaging plasmids: pMD.G and CMVdR8.74 (gift from the Naldini lab). Viral supernatant was harvested at 48 and 72 hours, filtered through a 0.45 μm syringe and frozen at −80°C or used fresh. Cell transductions were performed using a 1:4 dilution of lentiviral supernatant to media for 24 hours and culture media was subsequently changed.

ccRCC cell lines were transduced with lentivirus lentiCas9-Blast (Addgene Plasmid #52962) and selected with 4 μg/mL blasticidin for 10 days. Monoclonal lines were subsequently selected following limiting dilutions. Functional testing (Supplemental Figure 3C) was performed after transduction with lentivirus pXPR_011 (Supplemental Figure 3B, Addgene Plasmid #59702) and 7 days in culture.

Guide RNA sequences targeting PRMT1, PRMT3, PRMT4/CARM1, PRMT6, RPA3 and the human ROSA26 locus were selected from the Broad Insitutute’s ‘Brunello Library’ (https://portals.broadinstitute.org/gppx/crispick/public, Supplementary Table 2). Respective oligonucleotides were ordered from IDT and then cloned into the sgRNA delivery LRG plasmid (Addgene Plasmid #65656) using a BsmBI digestion.

Doxycycline inducible shRNA constructs targeting PRMT1 (shRNA target sequence: GTGTTCCAGTATCTCTGATTA) and Luciferase (non-targeting control, shRNA target sequence: CAAATCACAGAATCGTCGTAT) were obtained as a gift from the Structural Genomics Consortium. Protein coding sequences for PRMT1v1 (ENST00000391851.8) and PRMT1v2 (ENST00000454376.7) were ordered from GenScript (https://www.genscript.com) and PCR-cloned into pHIV-Luc-ZsGreen (Addgene plasmid #39196) using and EcoRI and XbaI digestion.

Mini-Turbo plasmid (pSTV6-N-miniTurbo-BirA) was obtained as a gift from Dr. Brian Raught’s lab. Open reading frame for PRMT1v1 (ENST00000391851.8) was cloned into the Mini-Turbo plasmid via gateway cloning using pDONR223 (gift from Dr. Brian Raught’s lab) and BP/LR combined cloning reagents (Gateway BP clonase II enzyme mix, Thermo #11789020 and LR Clonase II Enzyme Mix, Thermo, # 11791019).

All cloned plasmids were amplified in OneShot^TM^ Stbl3^TM^ E. Coli (ThermoFisher, #C737303) and prepared using the PureLink^TM^ Quick Plasmid Miniprep Kit (Invitrogen, #K210010). Plasmid identification was verified via Sanger sequencing.

### CRISPR/Cas9 Competition Assay & GFP drop out screening

Cas9 expressing cell lines were plated at ∼60% confluency in 6 well plates and transduced with their respective cloned-LRG plasmid. Fresh media was replaced on cells 48 hours later (day 0), the culture was passaged at 1:4 and a baseline GFP+ percentage was measured using BD LSR2 analytical flow cytometer. Cells were subsequently passaged every 4 days at a ratio of 1:4 and the percentage of GFP+ cells was measured at each split. Dropout values represent the fold decrease in GFP+ cells at each passage, relative to the GFP+ percentage on day 0.

### RNA Isolation,Sequencing and Pathway Analysis

RNA isolation was performed using the Qiashredder kit (Qiagen, #79654) and the RNeasy mini kit (Qiagen, #74104). RNA quality was assessed on an RNA 6000 Pico chip (Agilent Technologies) using the Agilent Bioanalyzer to determine sample RNA integrity number (RIN) and quantified by the Qubit RNA HS assay kit (Life Technologies). All samples had RIN values > 9.7. RNA was sent to Genome Quebec at McGill University for sequencing. NEB stranded mRNA libraries were constructed and paired-end (100bp) next generation sequencing was performed on a Illumina NovaSeq 6000 S4 system.

Raw reads were processed using TrimGalore v0.6.6 to remove adaptor sequences (via cutadapt v3.0) and to assess read quality (via fastqc v0.11.5). Reads then were mapped to the Genome Reference Consortium Human Build 38 patch release 12 (GRCh38.p12) reference genome using STAR v2.7.9a. Read counts per gene were subsequently obtained using the htseq-count command in the HTSeq v0.11.0 package alongside Gencode’s human genome annotation release 30. Differential gene expression analysis based on the negative binomial distribution was performed on treated vs untreated cells using the DESeq2 R package v1.36.0. Genes were considered differentially expressed if they were found with a minimum |log2(fold-change)|≥ 1 and FDR adjusted p-value ≤ 0.05.

Gene Ontology (GO) enrichment analysis was performed on differentially expressed lists using the PANTHER17.0 analysis tool^53^, which uses Fisher’s exact test and references the GO Ontology database for biological processes (DOI:10.5281/zenodo.6399963, released 2022-03-22)^54, 55^. Pathways were considered significant if they were found enriched at a minimum level of 2-fold above expectation and with a FDR adjusted p-value ≤ 0.05.

### Cell cycle/DNA content analysis

RCC243 cells were treated with 5 μM MS023 for the indicated time periods and then cells were trypsinized and stained using the Propidium Iodide Flow Cytometry Kit (Abcam, #ab139418, Abcam) based on the manufacturer’s instructions. A BD LSR2 analytical flow cytometer was used to acquire fluorescence data. All the flow cytometric data were analyzed using FlowJo software version 10.7.1 to quantify DNA content of individual cells and assess cell cycle status.

### Immunofluorescence and Focii Staining

Cells were grown on chamber slides and treated as indicated with MS023 or 0.1% DMSO. Following treatment, cells were washed three times in PBS and fixed in 4% paraformaldehyde for 15 min on ice. Cells were again washed 3 times with ice cold PBS and then permeabilized with two different blocking buffers: for Anti-phospho-Histone H2A.X (Ser139) - 5% BSA, 10% FBS, 0.25% Triton X-100, 1% fish skin gelatin in PBS; and for FANCD2 - 5% Normal Goat Serum, 0.3% Triton X-100 in PBS. Primary antibody was added in blocking buffer and incubated at 4°C overnight. Primary antibodies used: Anti-phospho-Histone H2A.X (Ser139) (Millipore, #05-636, 1:1000) and FANCD2 (CST, #16323, 1:100). Cells were washed 3 times in 0.25% Triton X-100 in PBS (PBS-T) and secondary antibodies: Goat anti-Rabbit IgG H+L -Alexa FluorTM 594 (Invitrogen #A11037, 1:400) or Goat anti-Mouse IgG (H+L) - Alexa FluorTM 488 (Invitrogen #A10667, 1:1000) was added in blocking buffer without 0.25% Triton X-100 for 1 hour at room temperature. Cells were again washed 3 times in 0.25% PBS-T and 1:1000

Hoechst was added in PBS for 1 min. A final wash in PBS-T was performed and coverslips were subsequently mounted using Mowiol. Images were collected on the Zeiss Axioimager Z1 wide field fluorescence microscope. Image processing and foci quantification was performed using Image J.

### BioID Sample Processing

Cells were transduced with Flag-miniTurbo-PRMT1v1 lentiviral supernatant for 48 hours before media was changed and puromycin selection applied. Cells were cultured for 1 week in the presence of puromycin and expression of mini-Turbo fusion protein was induced by addition of doxycyline (1 μg/mL) for 24 hours in surviving cells and confirmed via western blotting (Supplementary Figure 5). A total of 250 mg of Biotin (BioBasic ref# BB0078) was dissolved in 2.04 ml of NH_4_OH 28-30% (SIGMA ref# 221228) to produce a 500 mM solution. The solution was neutralized by gently adding 18 ml of 1N HCl to get a final stock solution at 50 mM. Biotin was then added to mini-Turbo expressing cells for 90 min to allow biotin tagging to proceed. Cells were harvested and frozen at −80°C. Pellets were lysed in modified RIPA (1% Triton X-100, 0.1% SDS, 50mM Tris-HCl pH 7.5, 1mM EDTA, 1mM EGTA, 150 mM NaCl, 0.5% sodium deoxycholate) supplemented with protease inhibitors and benzonase. Samples were incubated 1 hour at 4°C and sonicated prior to centrifugation at 15,000 x g for 30 minutes. Cleared lysates were incubated for 3 hours at 4°C and rinsed with 50 mM NH4HCO3 for 6 cycles. Trypsin (1 μg/sample; diluted in 50 mM NH4HCO3) was added to the beads and samples were incubated for 16 hours at 37°C. Additional trypsin (0.5 μg/sample) was added and samples were incubated for 2 hours at 37°C. Supernatant was collected, beads were rinsed with 50 mM NH4HCO3 eluates were pooled prior to the peptides being lyophilized in a vacuum centrifuge.

### Liquid chromatography – Mass Spectrometry (LC-MS)

Dried peptides were reconstituted in 0.1% HCOOH and injected on a loading column (C18 Acclaim PepMapTM 468 100, 75μM x 2cm, 3μm, 100Å) prior to separation on an analytical column (C18 Acclaim PepMapTM 469 RSLC, 75μm x 50cm, 2μm, 100Å) by HPLC over a reversed-phase gradient (120-minute gradient, 5-30% CH_3_CN in 0.1% HCOOH) at 225nL/min on an EASY-nLC1200 pump in-line with a Q-Exactive HF (Thermo Scientific) mass spectrometer operated in positive ESI mode. An MS1 ion scan was performed at 60,000 fwhm followed by MS/MS scans (HCD, 15,000 fwhm) of up to 20 parent ions (minimum activation of 1000). Fragmented ions were added to a dynamic exclusion list (10 ppm) for 5 seconds. For peptide and protein identification, raw files (.raw) were converted to .mzML format with Proteowizard (v3.0.19311) and searched using X!Tandem (v2013.06.15.1) and Comet (v2014.02.rev.2) search engines against the human proteome RefSeqV104 database (36,113 entries). Parent ion mass tolerance was set at 15 ppm and an MS/MS fragment ion mass tolerance at 0.4 Da. Up to two missed cleavages was allowed. No fixed modification was set. Deamidation (NQ), oxidation (M), acetylation (protein N-term) were set as variable modifications. Search results were further processed using the trans-proteomic pipeline (TPP v4.7) using iProphet. Proteins were identified with an iProphet cut-off of 0.9 and at least two unique peptides. Putative proximity interactors were identified using Significance Analysis of INTeractome (SAINT)^56^ comparing Flag-miniTurbo-only samples to Flag-miniTurbo-PRMT1 samples using a Bayesian false discovery rate (BFDR) cut-off of ≤0.01 (1%). For visualization representation, the protein names of interactors identified in both RCC243 and 786-0 cells (n=41) were imported into Cytoscape 3.9.1 and enrichment analysis was carried out using the STRING Enrichment app using the categories GO biological process, GO molecular function and COMPARTMENTS.

### Immunoprecipitation

RCC243 cells were seeded in 10 cm plates and allowed to reach ∼60% confluence. Cells were treated with 5 μM MS023 or 0.1% DMSO for three days. Cells were collected and lysed in 1 mL ice cold RIPA buffer supplemented with HaltTM Protease Inhibitor Cocktail (Thermo # 78425) and sonicated 2 times for 10 sec @ 50% output followed by a 10 min incubation on ice. After cell lysis and sonication, tubes were centrifuged, and supernatant was transferred into a new Eppendorf tube.

Protein A Sepharose beads (Abcam, #ab193256) were washed with 1 mL of RIPA buffer and added to each sample tube to facilitate pre-clearing of the samples. Beads and samples were shaken for 1 hour at 4°C on a nutator and the supernatant was transferred to a new Eppendorf tube. BCLAF1 antibody (Bethyl A300-608A) or mouse IgG was added and incubated at 4°C overnight with continuous shaking on a nutator. New protein A beads were washed in RIPA and added to each sample the next day and incubated for 3 hours at 4°C on a nutator. Samples were then centrifuged for 1 min at 1000xg and the beads collected and washed 3 times in RIPA. Beads were resuspended in 50 μL SDS loading buffer, boiled at 95°C and run on a Western blot (as described in section 2.5.4) probed with BCLAF1 antibody (Bethyl A300-608A, 1:1000) and Asymmetric Di-Methyl Arginine ASYM25 (Sigma, #09-814, 1:1000).

### Cytoplasmic/Nuclear RNA Fragmentation

Cells were typsinized and pelleted by centrifugation at 300 x *g* for 5 min. The plasma membrane was then lysed with 175 µl of cooled RLN buffer (50 mM Tris-HCl, pH 8.0, 140 mM NaCl, 1.5 mM MgCl_2_, 0.5% (v/v) Nonidet P-40 (1.06g/ml) for 5 minutes on ice. Lysate was then centrifuged at 4°C for 2 minutes at 300 x *g*. The supernatant, containing the cytoplasmic RNA, was transferred to a new tube and the remaining pellet contained the nuclear fraction. RNA was then extracted from both the cytoplasmic and nuclear fractions using the Qiagen RNeasy mini kit. The purity of cytoplasmic and nuclear fractions were assessed for the expression of the nuclear enriched MALAT1 RNA via quantitative reverse transcriptase (qRT)-PCR.

### qRT-PCR

Synthesis of cDNA was performed on 1 µg of DNAse-free RNA using the SuperScript VILO cDNA Synthesis Kit (Invitrogen, Thermo Fisher). Power SYBR Green PCR Master Mix (Invitrogen, Thermo Fisher) was used to perform qPCR on 10 ng of cDNA for each sample (in triplicate) using primers specific to each transcript. Samples were loaded into a Bio-Rad CFX96 real-time PCR detection system following the manufacturer’s protocol. Post-spliced transcripts (exon-exon spanning primers) and pre-spliced transcripts (exon-intron spanning primers) were normalized to ACTB expression levels within the same samples. The expression of each gene of interest was normalized to ACTB and then the relative amounts of expression were calculated by the delta delta Ct method. The primers used were as follows:

MALAT1 Forward: 5′- GACGGAGGTTGAGATGAAGC-3′, MALAT1 Reverse: 5′-

ATTCGGGGCTCTGTAGTCCT- 3′, ACTB Forward: 5′-AGACCTGTACGCCAACACAG-3′, ACTB

Reverse: 5′- GGAGCAATGATCTTGATCTTCA-3′ FANCD2_Exon36_Forward: 5′-

CCCAGAACTGATCAACTCTCCT-3′, FANCD2_Exon37_Reverse: 5′-

CCATCATCACACGGAAGAAA-3′, FANCD2_Intron37_Reverse: 5′-

ACAGGTGTGTGCCACCGTG-3′, BRCA2_Exon18_Forward: 5′- CCTGATGCCTGTACACCTCTT-

3′, BRCA2_Exon19_Reverse: 5′- GCAGGCCGAGTCACTGTTAGC-3′, BRCA2_Intron18_Reverse:

5′- TACATCTAAGAAATTGAGCATCCT-3′, Rad51_Exon6_Forward: 5′-

TGAGGGTACCTTTAGGCCAGA-3′, Rad51_Exon7_Reverse: 5′-

CACTGCCAGAGAGACCATACC-3′, Rad51_Intron6_Reverse: 5′-

AGAGACATTCTTCGGCCAAACT-3′ and FANCA_Exon2_Forward: 5’-

TCCTGAAAGGGCACAGAAATTA-3’, FANCA_Exon2/3_Reverse: 5’-

GGGCCTTCTACCTCAAGCAAA-3’, FANCA_Intron2_Reverse: 5’-

CCAGCTTCCTCTTACCTCAAG-3’, RAD51 AP1_Exon2_Forward: 5’-

CCAGTCAATTACTCACAGTTTGAC-3’, RAD51 AP1_Exon3_Reverse: 5’-

TAACTCCTTTGGTGCTGTTCT-3’, RAD51 AP1_Intron2_Reverse: 5’-TCCGAGGAAA

TGAGTTTCCAA-3’.

### In vivo tumor studies and drug treatments

ccRCC samples were obtained from the University Health Network (UHN) and the Cooperative Health Tissue Network from patients providing written consent under UHN Research Ethics Board approval, protocol #09-0828-T. PDX models were generated by implanting patient tissue under the renal capsule of 6-8 week old male NOD/SCID/IL2Rγ−/− (NSG) mice and allowed to expand until humane endpoint was reached. All animal studies were approved by the University Health Network Animal Care Committee under protocol #4896. Once tumors were established in the renal capsule, tissue was harvested and finely minced using a scalpel, then incubated in 1× Collagenase/Hyaluronidase and 125 Units/mL DNase (Stem Cell Technologies) with frequent pipetting at 37 °C for two hours. Red blood cells were lysed with ammonium-chloride/potassium (ACK) lysing buffer (Gibco) and cell clumps were filtered using 70 μm nylon mesh. Dissociated cells were stained with trypan blue, viable cells were counted for re-implantation into mice. 1-5 million viable cells were injected in 100 μL of phosphate-buffered saline (PBS):Matrigel (1:1) in the subcutaneous left flank of 6-8 week old NSG mice. Tumors were allowed to grow to 100-200 mm^3^ then randomized into each treatment group of 5 mice per arm. MS023 was administered 3 days sequentially at 80 mg/kg intraperitoneally, with 4 days off for a period of three weeks or until vehicle control reached humane endpoint. Tumor measurements were collected twice a week for the duration of the experiment. Cell lines were injected as described above. For inducible cell line studies, doxycycline treatment was initiated once tumors reached between 100-200 mm^3^ and was delivered into the drinking water at 1 mg/ml. Water was changed twice a week until endpoint. Tumors were excised at endpoint and tumor fragments were snap frozen in liquid nitrogen and stored at −80 °C for further analysis.

### Comet Assay

Cells were plated in 10 cm culture dishes at ∼60% confluence and incubated for 24 hours with 5 μM MS023, 2 μM NU7441 (DNA-PK inhibitor, gift from Dr. Shane Harding) or 0.1% DMSO. Cells were then exposed to 10 Gy ionizing radiation (IR) using a Cs137 irradiator and were collected at indicated time points post (IR) exposure. Single cell electrophoresis was carried out in the CometAssay® Electrophoresis System II (Trevigen, #4250-050-ES) in accordance with the manufacturer’s protocol for neutral conditions (double stranded break quantification). Microscopy was carried out on the Zeiss Axioimager Z1 wide field fluorescence microscope and comet tail moments were calculated in ImageJ with the OpenComet plugin v1.3.

### Statistics and Reproducibility

Statistical analysis was performed using GraphPad Prism 9.3.1 using statistical tests described in figure legends.

## Acknowledgements

We thank the patients who graciously donated their samples for the study. We would like to thank Dr. Shane Harding for advice and reagents relating to DNA damage assays. We would also like to thank Dr. Jian Jin and Yudao Shen for the synthesis of MS023 used for *in vivo* studies. This research is supported by the Canadian Cancer Society (grant #23145) and the Ontario Institute for Cancer Research (IA-016). The Structural Genomics Consortium is a registered charity (no: 1097737) that receives funds from Bayer AG, Boehringer Ingelheim, Bristol Myers Squibb, Genentech, Genome Canada through Ontario Genomics Institute [OGI-196], EU/EFPIA/OICR/McGill/KTH/Diamond Innovative Medicines Initiative 2 Joint Undertaking [EUbOPEN grant 875510], Janssen, Merck KGaA (aka EMD in Canada and US), Pfizer and Takeda.

## Supplemental Figure Legends

**Supplemental Figure 1**

Epiprobe chemical screen results across all cell lines. Data are presented as the mean fluorescence ± SEM and *p* values are calculated by 1-way ANOVA with Dunnett’s multiple comparison relative to DMSO control group. ***, p <0.001, ****, p<0.0001.

**Supplemental Figure 2**

Dose response curves across cell lines RCC407, RCC364, RCC323, RCC243 and RCC22, for **a,** GSKJ4 vs inactive control GSKJ5. **b,** (+)JQ1 vs inactive control (-)JQ1. **c,** UNC1999 vs inactive control UNC2400. **d,** GSK591 vs inactive control SGC2096. **e,** MS023 vs inactive control MS094. **f,** PF11

**Supplemental Figure 3**

**a,** Western blot analysis of Cas9 expression in RCC243_Cas9_, 786-0_Cas9_ and RCC364_Cas9_ following transduction with lentiviral delivered humanized Cas9 gene and serial dilutions to establish monoclonal lines. **b,** Functional testing, lentiviral genetic construct that expresses both the green fluorescence protein (GFP) and the guide (g-)RNA targeting GFP. **c,** Analytical flow cytometry analysis of GFP expression in parental and Cas9-expressing lines 5 days after genetic cassette in (b) introduced *via* lentiviral transduction. The gating strategy to select for viable, non-doublet cells is shown to the right.

**Supplemental Figure 4**

RCC243 cells were treated with 5 μM MS023, 2 μM DNA-PK-inhibitor NU7441 or control conditions and then exposed to 10 Gy of irradiation and allowed to recover for the indicated time. Cells were then lysed and analyzed by neutral comet assay. Comet tail moment averages are shown with error bars representing the SEM of three independent experiments. p-values are calculated by 2-way ANOVA with repeated measures and Sidak’s multiple comparisons test. n =5 to 10 mice/group. *, p<0.05; ***, p<0.001. Representative analysis of images from MS023 treated time course pictured above graphs.

**Supplemental Figure 5**

**a,** RCC243 cells were transduced with PRMT1-miniTurbo fusion (pSTV6-Nterm-PRMT1) or miniTurbo alone (pSTV6-N) and 1 μg/mL doxycycline was added for 24 hours to induce protein expression. 50 μM biotin was then added for the indicated times. Western blot analysis of protein expression *via* FLAG-tag detection and biotinylation levels *via* streptavidin detection are shown. **b,** 786-0 cells were transduced with PRMT1-miniTurbo fusion (pSTV6-Nterm-PRMT1) or miniTurbo alone (pSTV6-N) and 1 μg/mL doxycycline was added for 24 hours to induce protein expression. 50 μM biotin was added for 90 min. Western blot analysis of protein expression *via* FLAG-tag detection and biotinylation levels *via* streptavidin detection are shown.

**Supplemental Figure 6**

MALAT1 lncRNA levels in the nuclear vs cytoplasmic fractions for samples in Figure 7F as assessed by qRT-PCR. MALAT1 is largely absent from cytoplasmic fractions relative to nuclear compartments.

**Supplemental Figure 7**

**a,** Dosing and tissue collection schedule for PK study. **b,** Mass spectrometry serum measurements of MS023 post Day 3 injection. **c,** Mass spectrometry tissue measurements of MS023 in tumor tissue, kidney and liver post Day 3.

