## Supplemental Tables for "PRMT1 is a critical dependency in clear cell renal cell carcinoma through its role in post-transcriptional regulation of DNA damage response genes"

### Supplementary Table 1 - Epiprobe chemicals

| Protein Family | Target | Probe | PubMed ID |
|---|---|---|---|
| Methyltransferase | SUV20H1/H2 | A-196 | 28114273 |
| Methyltransferase | EHMT2, EHMT1 | A-366 | 24900801 |
| Methyltransferase | SMYD2 | BAY-598 | 27075367 |
| Bromodomain | BAZ2A/2B | BAZ2-ICR | 25719566 |
| Bromodomain | BRD9, BRD7 | BI-9564 | 26914985 |
| Bromodomain | BAZ2A/2B | GSK2801 | 25799074 |
| Methyltransferase | EZH2 | GSK343 | 24900432 |
| PAD | PAD-4 | GSK484 | 25622091 |
| Methyltransferase | PRMT5 | GSK591 | 26985292 |
| KDM | KDM6B, KDM6A | GSKJ4 | 22842901 |
| KDM | KDM1A | GSKLSD1 | 26175415 |
| Bromodomain | BRD9 | I-BRD9 | 25856009 |
| Bromodomain | CREBBP P300 | I-CBP112 | 26552700 |
| PHD | pan-2-OG | IOX1 | 24504543 |
| Bromodomain | BRD2-4 | JQ-1 | 20871596 |
| Bromodomain | BRD9, BRD7 | LP99 | 25864491 |
| Methyltransferase | Type I PRMT | MS023 | 26598975 |
| Methyltransferase | PRMT4, PRMT6 | MS049 | 27584694 |
| Bromodomain | BRPF1-3 | NI-57 | 28714688 |
| Bromodomain | BRPF1-3 | OF-1 | 21804994 |
| Bromodomain | BRD2-4 | PFI-1 | 23095041 |
| Methyltransferase | SETD7 | PFI-2 | 25136132 |
| Bromodomain | SMARCA2,4 | PFI-3 | 26139243 |
| Bromodomain | BRPF1B | PFI-4 | 28849908 |
| Methyltransferase | DOT1L | SGC0946 | 23250418 |
| Methyltransferase | PRMT3 | SGC707 | 25728001 |
| Bromodomain | CREBBP P300 | SGC-CBP30 | 24946055 |
| Methyltransferase | EHMT2, EHMT1 | UNC0638 | 21743462 |
| Kme | L3MBTL3 | UNC1215 | 23292653 |
| Methyltransferase | EZH2 | UNC1999 | 23614352 |
| Bromodomain | Pan-Bromodomain | Bromosporine | 27757418 |
| Bromodomain | CECR2 | NVS-CECR2-1 | - |
| HDAC | HDAC | LAQ824 | 12816865 |
| HDAC | HDAC | CI-994 | 17455259 |
| Methyltransferase | EHMT2, EHMT1 | UNC0642 | 24102134 |
| WD40 repeats | WD40 (WDR5) | OICR-9429 | 26167872 |

**Supplementary Table 2 - sgRNA's**

| sgRNA name | sgRNA Target Sequence | PAM Sequence | Target Transcript | Exon Number (Relative to transcript) | Strand | On-Target Efficacy Score (Azimuth 2.0) | Position of Base After Cut (1-based) |
|---|---|---|---|---|---|---|---|
| PRMT1_e4 | AAAGCCAACAAGTTAGACCA | CGG | NM_198318.4 (PRMT1v1) | 4 | sense | 0.7305 | 49682255 |
| PRMT1_e6 | GATGGCCGTCACATACAGCG | TGG | NM_198318.4 (PRMT1v1) | 6 | antisense | 0.7528 | 49684791 |
| PRMT1_e7 | GGGTCCACGACATCCACTAG | GGG | NM_198318.4 (PRMT1v1) | 7 | antisense | 0.7112 | 49684987 |
| PRMT1_e2 | GTGGATGCCAAAGTGTGCGT | AGG | NM_198318.4 (PRMT1v1) | 2 | antisense | 0.6442 | 49680569 |
| PRMT3e8 | CCAACATCCAAAACTACCTA | GGG | NM_001145167.1 | 8 | antisense | 0.6114 | 20407911 |
| PRMT3e4 | GAATTCATGTACTCAACTGT | AGG | NM_001145167.1 | 4 | antisense | 0.5647 | 20392906 |
| PRMT3e6 | GTCATCTACTAGTGTCATTG | CGG | NM_001145167.1 | 6 | sense | 0.6283 | 20397642 |
| PRMT3e9 | TAGATGTTATCATATCTGAG | TGG | NM_001145167.1 | 9 | sense | 0.7266 | 20426857 |
| PRMT6e1.1 | CGTTCCCGCTTAGTCCTCCG | GGG | NM_018137.2 | 1 | antisense | 0.7238 | 107056827 |
| PRMT6e1.2 | CTCGGACGTTTCGGTCCACG | AGG | NM_018137.2 | 1 | sense | 0.6903 | 107056885 |
| PRMT6e1.3 | GCCCATCCACTCGCTCACGA | TGG | NM_018137.2 | 1 | antisense | 0.6718 | 107057173 |
| PRMT6e1.4 | GTGGCCCATGAGACAGCGCG | TGG | NM_018137.2 | 1 | antisense | 0.6809 | 107057398 |
| CARM1e7 | TAGAGCTGTTCATCCGTGAA | GGG | NM_199141.1 | 7 | antisense | 0.6214 | 10916451 |
| CARM1e1 | TCGCGTCGCCGATGGTGAGG | AGG | NM_199141.1 | 1 | antisense | 0.6159 | 10871816 |
| CARM1e5 | TGGAGCACGGAAAATCTACG | CGG | NM_199141.1 | 5 | sense | 0.8313 | 10912257 |
| CARM1e6 | TTGAAGAGCATGTAGCCCAT | GGG | NM_199141.1 | 6 | antisense | 0.631 | 10913985 |
| RPA3_e5.1 | GATGAATTGAGCTAGCATGC | CGG | NM_002947.3 | 5 | antisense | 0.6306 | 7640373 |
| RPA3_e7 | GGTTGGAAGAGTAACCGCCA | AGG | NM_002947.3 | 7 | sense | 0.7045 | 7637924 |
| RPA3_e6 | TACGGGTTCCATCAACTCGA | TGG | NM_002947.3 | 6 | antisense | 0.6307 | 7639084 |
| RPA3_e5.2 | TGGACATGATGGACTTGCCC | AGG | NM_002947.3 | 5 | sense | 0.6221 | 7640398 |
| Rosa26 g1 (THUMPD3-AS1) | ACCTCTCTGTGCTTACGCAA | NGG | NR_132780.1 | 4 | antisense | 0.6255 | 9390454 |
| Rosa26 g2 (THUMPD3-AS1) | GTTGCTTTATCACTCAACAG | NGG | NR_132780.1 | 4 | antisense | 0.7707 | 9389195 |
| Rosa26 g3 (THUMPD3-AS1) | GGAATGATCTTGAATAGTGT | NGG | NR_132780.1 | 4 | sense | 0.5791 | 9389474 |
| Rosa26 g4 (THUMPD3-AS1) | AAGCAGTAGTCAAGATCACC | NGG | NR_132780.1 | 2 | antisense | 0.5735 | 9395543 |
