## Supplemental Figures for "PRMT1 is a critical dependency in clear cell renal cell carcinoma through its role in post-transcriptional regulation of DNA damage response genes"

### Supplemental Figure 1

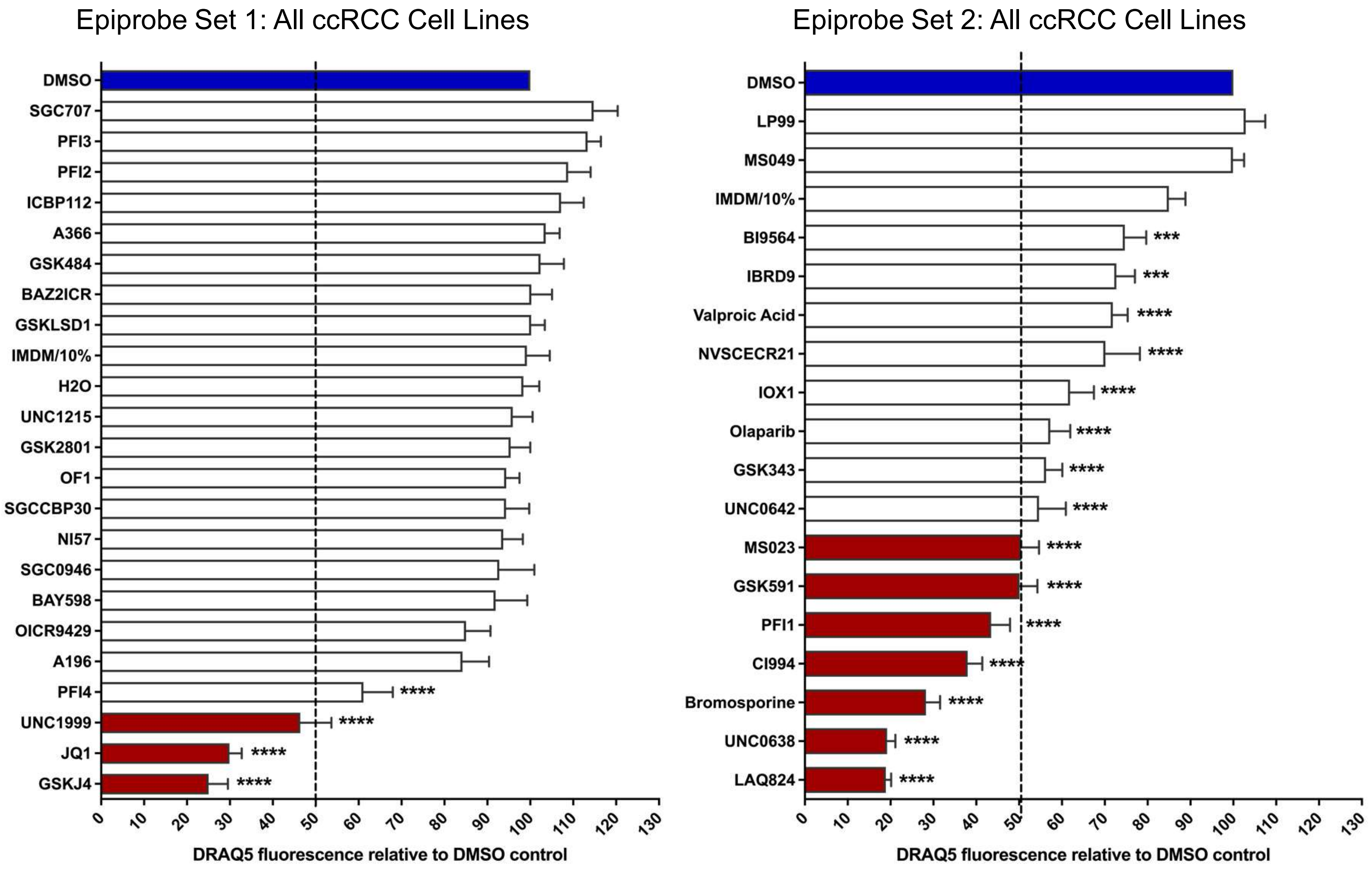

Supplemental Figure 2

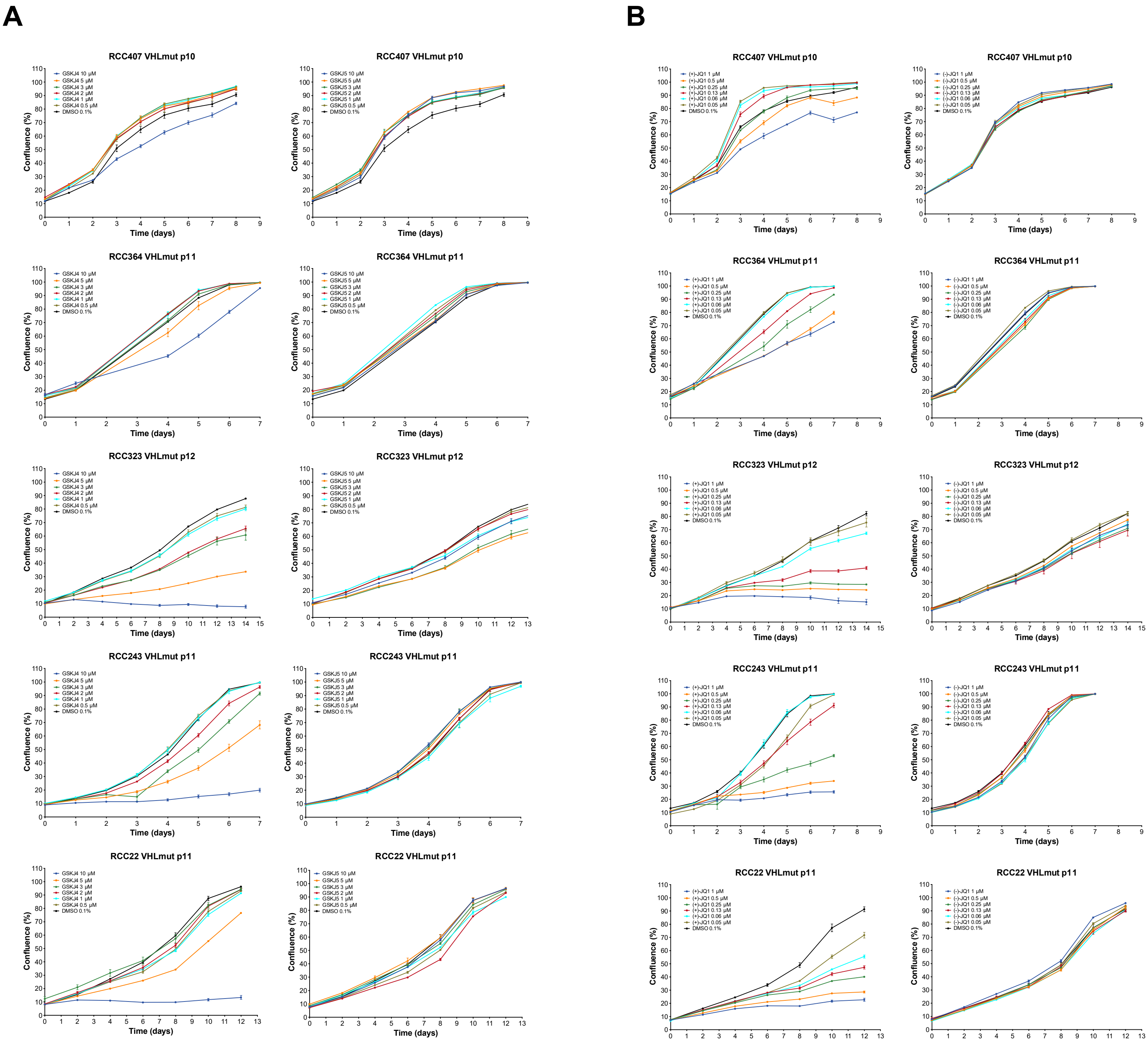

C

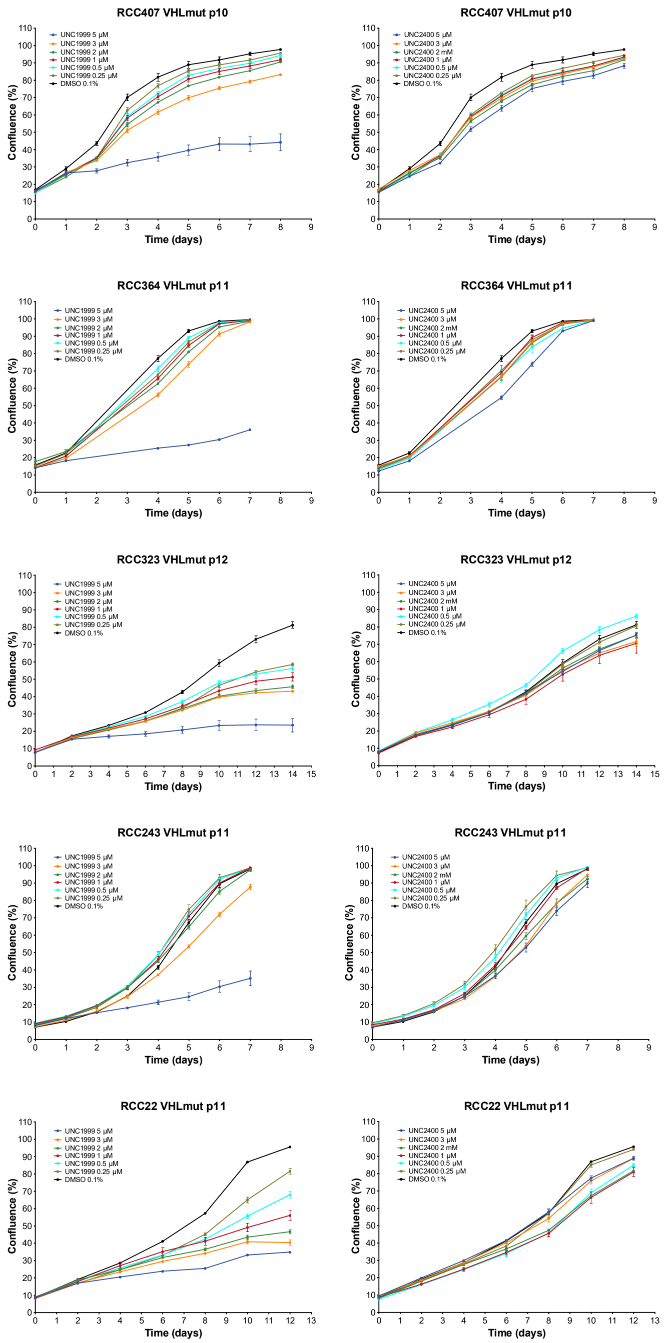
RCC407 VHLmut p10
RCC364 VHLmut p11
RCC323 VHLmut p12
RCC243 VHLmut p11
RCC22 VHLmut p11
UNC1999 5 µM
UNC1999 3 µM
UNC1999 2 µM
UNC1999 1 µM
UNC1999 0.5 µM
UNC1999 0.25 µM
UNC2400 5 µM
UNC2400 3 µM
UNC2400 2 mM
UNC2400 1 µM
UNC2400 0.5 µM
UNC2400 0.25 µM
DMSO 0.1%
Confluence (%)
Time (days)

D

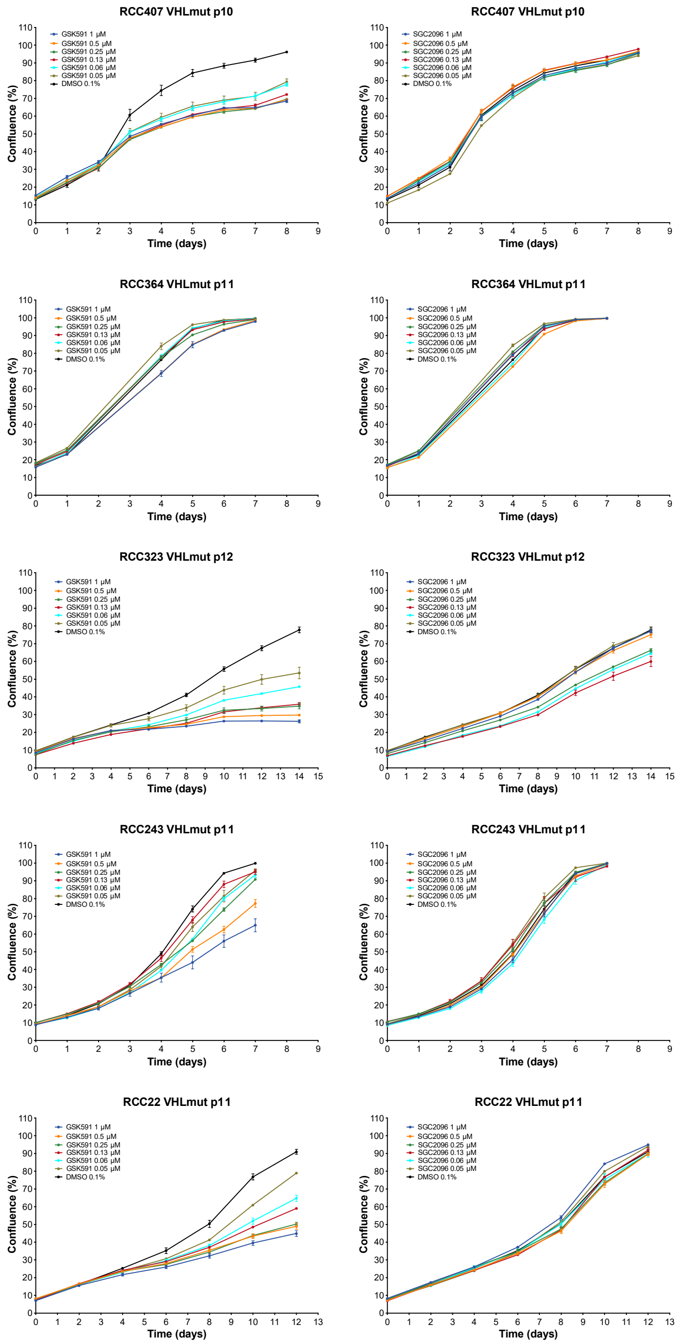
RCC407 VHLmut p10
RCC364 VHLmut p11
RCC323 VHLmut p12
RCC243 VHLmut p11
RCC22 VHLmut p11
GSK591 1 µM
GSK591 0.5 µM
GSK591 0.25 µM
GSK591 0.13 µM
GSK591 0.06 µM
GSK591 0.05 µM
SGC2096 1 µM
SGC2096 0.5 µM
SGC2096 0.25 µM
SGC2096 0.13 µM
SGC2096 0.06 µM
SGC2096 0.05 µM
DMSO 0.1%
Confluence (%)
Time (days)

E

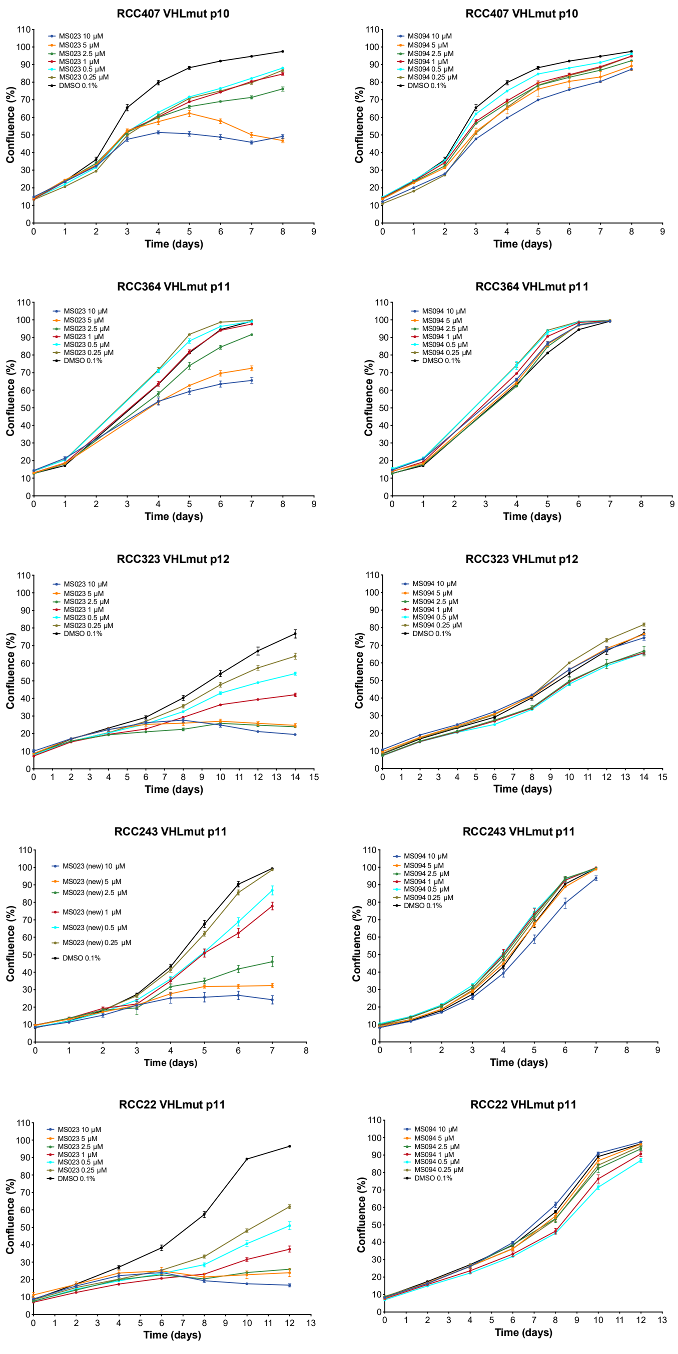
RCC407 VHLmut p10
MS023 10 µM
MS023 5 µM
MS023 2.5 µM
MS023 1 µM
MS023 0.5 µM
MS023 0.25 µM
DMSO 0.1%
Confluence (%)
Time (days)
RCC407 VHLmut p10
MS094 10 µM
MS094 5 µM
MS094 2.5 µM
MS094 1 µM
MS094 0.5 µM
MS094 0.25 µM
DMSO 0.1%
Confluence (%)
Time (days)
RCC364 VHLmut p11
MS023 10 µM
MS023 5 µM
MS023 2.5 µM
MS023 1 µM
MS023 0.5 µM
MS023 0.25 µM
DMSO 0.1%
Confluence (%)
Time (days)
RCC364 VHLmut p11
MS094 10 µM
MS094 5 µM
MS094 2.5 µM
MS094 1 µM
MS094 0.5 µM
MS094 0.25 µM
DMSO 0.1%
Confluence (%)
Time (days)
RCC323 VHLmut p12
MS023 10 µM
MS023 5 µM
MS023 2.5 µM
MS023 1 µM
MS023 0.5 µM
MS023 0.25 µM
DMSO 0.1%
Confluence (%)
Time (days)
RCC323 VHLmut p12
MS094 10 µM
MS094 5 µM
MS094 2.5 µM
MS094 1 µM
MS094 0.5 µM
MS094 0.25 µM
DMSO 0.1%
Confluence (%)
Time (days)
RCC243 VHLmut p11
MS023 (new) 10 µM
MS023 (new) 5 µM
MS023 (new) 2.5 µM
MS023 (new) 1 µM
MS023 (new) 0.5 µM
MS023 (new) 0.25 µM
DMSO 0.1%
Confluence (%)
Time (days)
RCC243 VHLmut p11
MS094 10 µM
MS094 5 µM
MS094 2.5 µM
MS094 1 µM
MS094 0.5 µM
MS094 0.25 µM
DMSO 0.1%
Confluence (%)
Time (days)
RCC22 VHLmut p11
MS023 10 µM
MS023 5 µM
MS023 2.5 µM
MS023 1 µM
MS023 0.5 µM
MS023 0.25 µM
DMSO 0.1%
Confluence (%)
Time (days)
RCC22 VHLmut p11
MS094 10 µM
MS094 5 µM
MS094 2.5 µM
MS094 1 µM
MS094 0.5 µM
MS094 0.25 µM
DMSO 0.1%
Confluence (%)
Time (days)

F

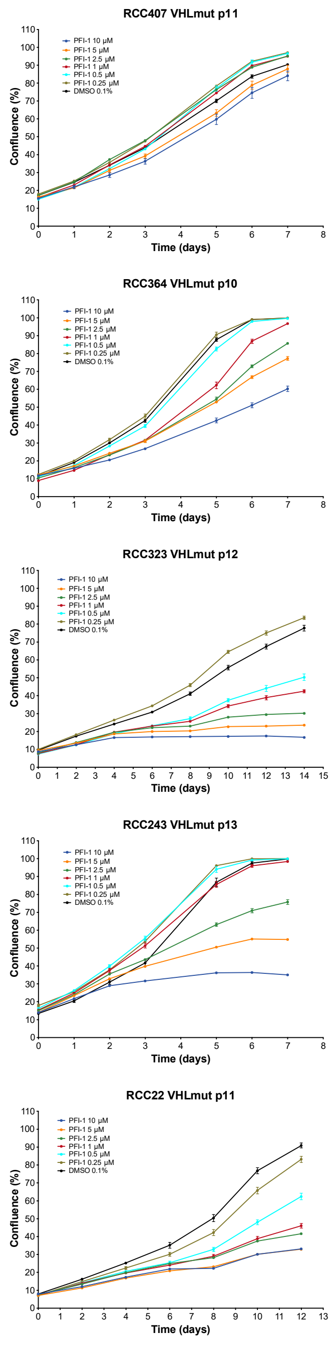
RCC407 VHLmut p11
PFI-1 10 µM
PFI-1 5 µM
PFI-1 2.5 µM
PFI-1 1 µM
PFI-1 0.5 µM
PFI-1 0.25 µM
DMSO 0.1%
Confluence (%)
Time (days)
RCC364 VHLmut p10
PFI-1 10 µM
PFI-1 5 µM
PFI-1 2.5 µM
PFI-1 1 µM
PFI-1 0.5 µM
PFI-1 0.25 µM
DMSO 0.1%
Confluence (%)
Time (days)
RCC323 VHLmut p12
PFI-1 10 µM
PFI-1 5 µM
PFI-1 2.5 µM
PFI-1 1 µM
PFI-1 0.5 µM
PFI-1 0.25 µM
DMSO 0.1%
Confluence (%)
Time (days)
RCC243 VHLmut p13
PFI-1 10 µM
PFI-1 5 µM
PFI-1 2.5 µM
PFI-1 1 µM
PFI-1 0.5 µM
PFI-1 0.25 µM
DMSO 0.1%
Confluence (%)
Time (days)
RCC22 VHLmut p11
PFI-1 10 µM
PFI-1 5 µM
PFI-1 2.5 µM
PFI-1 1 µM
PFI-1 0.5 µM
PFI-1 0.25 µM
DMSO 0.1%
Confluence (%)
Time (days)

Supplemental Figure 3

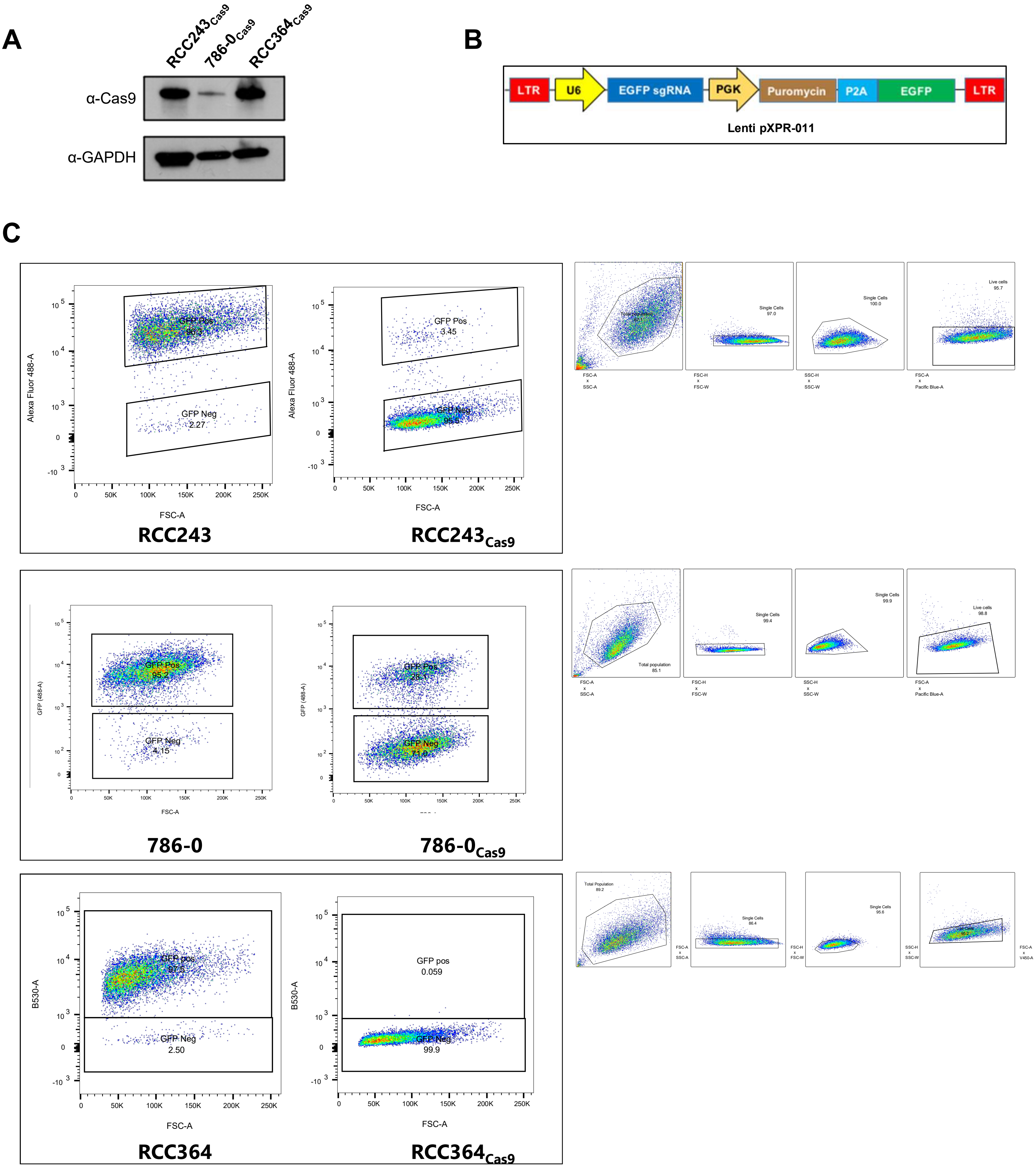

Supplemental Figure 4

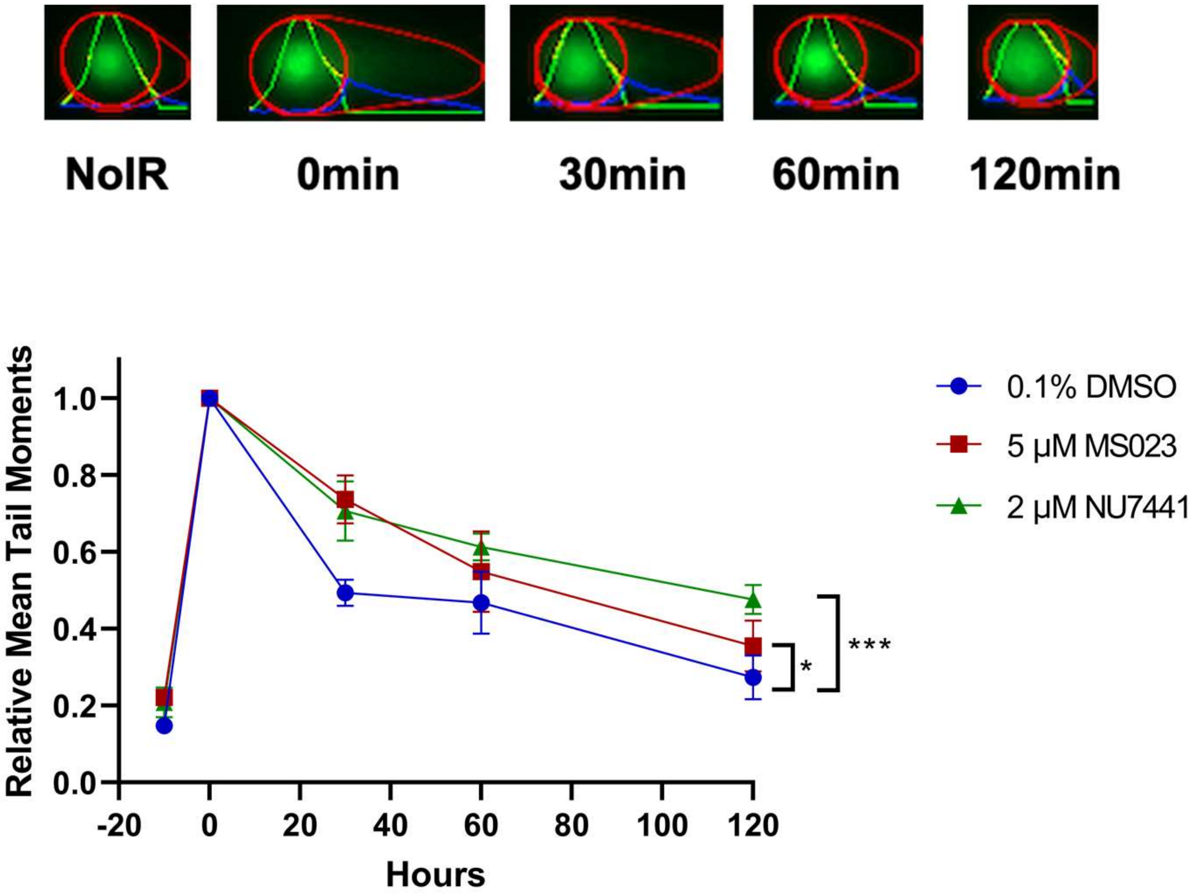

Supplemental Figure 5

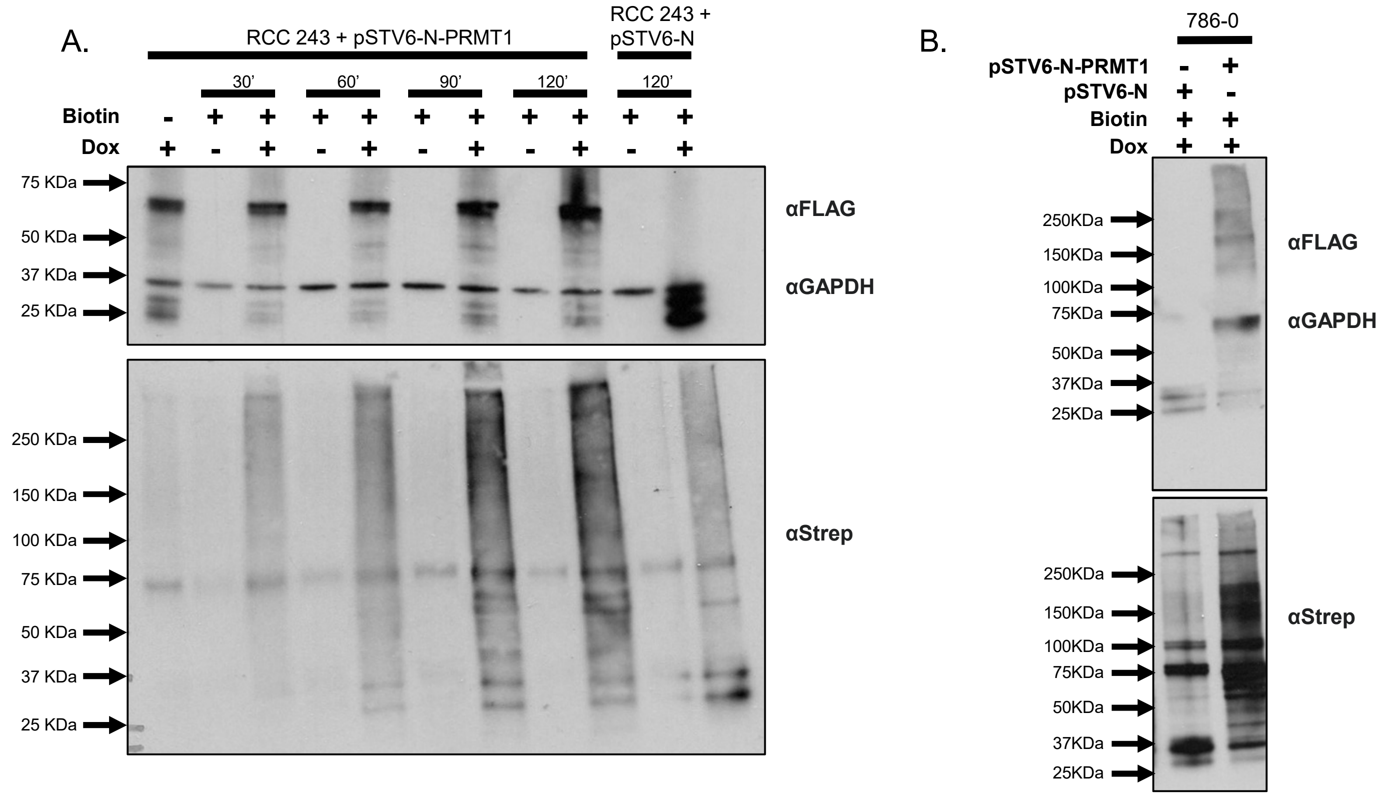

Supplemental Figure 6

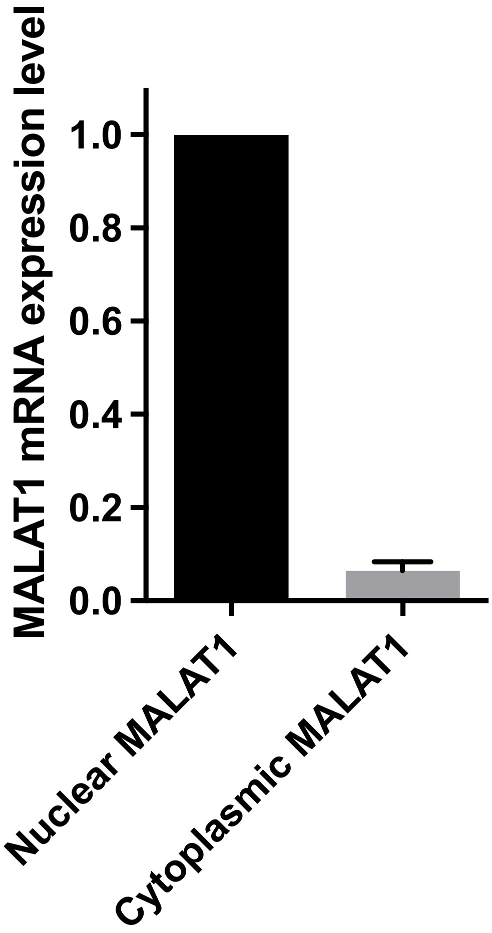

Supplemental Figure 7

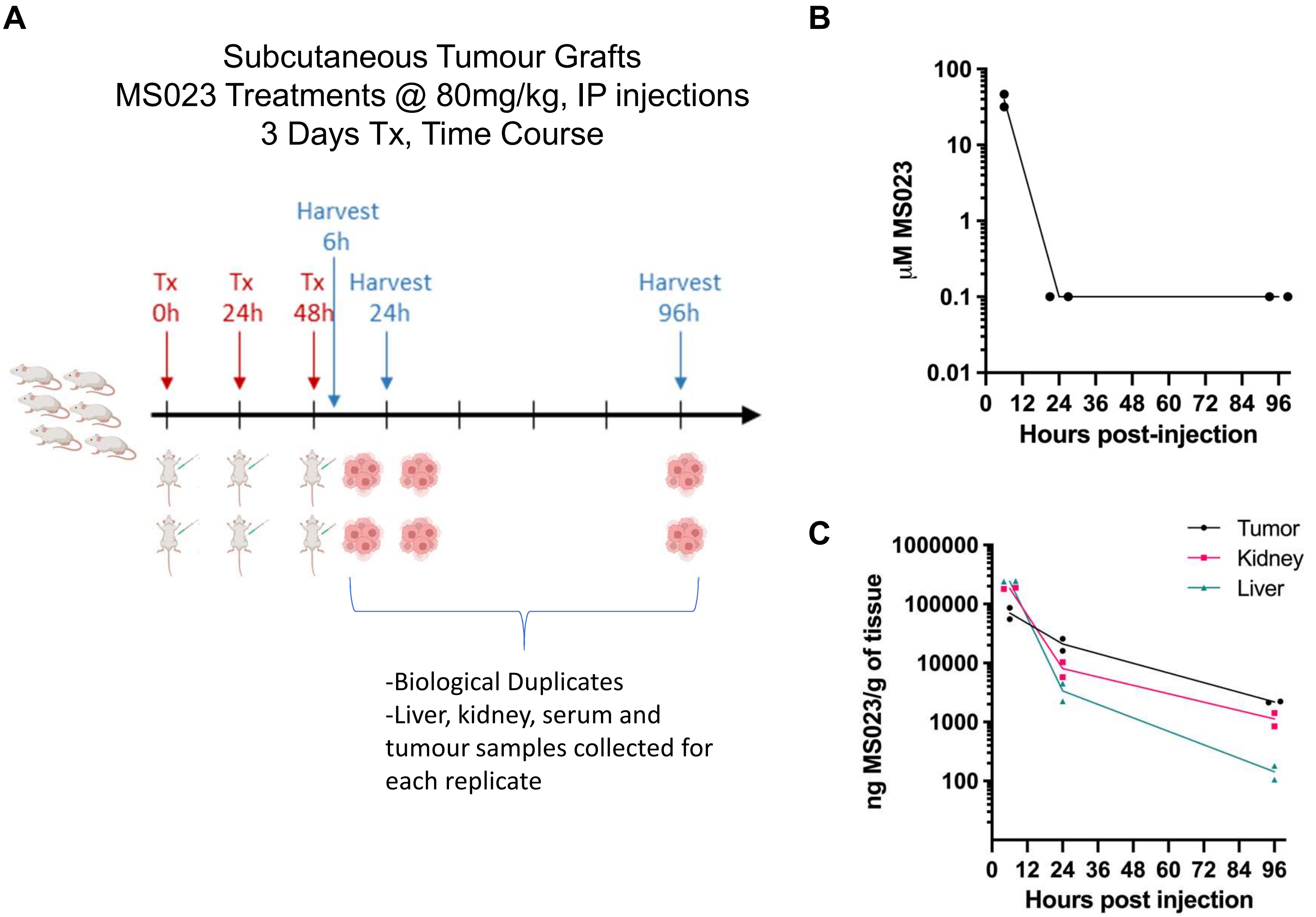
